# Synaptic alterations associated with disrupted sensory encoding in a mouse model of tauopathy

**DOI:** 10.1101/2023.05.24.542163

**Authors:** Soraya Meftah, Annalisa Cavallini, Tracey K. Murray, Lukasz Jankowski, Suchira Bose, Michael C. Ashby, Jonathan T. Brown, Jonathan Witton

## Abstract

Synapse loss is currently the best biological correlate of cognitive decline in Alzheimer’s disease and other tauopathies. Synapses seem to be highly vulnerable to tau-mediated disruption in neurodegenerative tauopathies. However, it is unclear how and when this leads to alterations in function in relation to the progression of tauopathy and neurodegeneration. We used the well-characterised rTg4510 mouse model of tauopathy, at 5-6 months and 7-8 months of age, to study the functional impact of cortical synapse loss. The earlier age was used as a model of prodromal tauopathy, with the later age corresponding to more advanced tau pathology and presumed progression of neurodegeneration. Analysis of synaptic protein expression in the somatosensory cortex showed significant reductions in synaptic proteins and NMDA and AMPA receptor subunit expression in rTg4510 mice. Surprisingly, *in vitro* whole-cell patch clamp electrophysiology from putative pyramidal neurons in layer 2/3 of the somatosensory cortex suggested no functional alterations in layer 4 to layer 2/3 synaptic transmission at 5-6 months. From these same neurons, however, there were alterations in dendritic structure, with increased branching seen proximal to the soma in rTg4510 neurons. Therefore, *in vivo* whole-cell patch clamp recordings were utilised to investigate synaptic function and integration in putative pyramidal neurons in layer 2/3 of the somatosensory cortex. These recordings revealed a significant increase in the peak response to synaptically-driven sensory stimulation-evoked activity and a loss of temporal fidelity of the evoked signal to the input stimulus in rTg4510 neurons. Together, these data suggest that loss of synapses, changes in receptor expression, and dendritic restructuring may lead to alterations in synaptic integration at a network level. Understanding of these compensatory processes could identify targets to help delay symptomatic onset within dementia.

**Abbreviated summary:** Meftah et al. report alterations to synaptic and dendrite properties in the rTg4510 mouse model of tauopathy associated with disrupted synaptic integration *in vivo*. Therefore, disrupted synaptic and network integration may be early markers of synapse loss in neurodegenerative tauopathies.

## Introduction

Tauopathies are a group of progressive neurodegenerative diseases, termed such due to their association with abnormal intracellular accumulations of tau protein.^1,2^ Cognitive decline in these diseases best correlates with synaptic degeneration, highlighting the synapse as a vulnerable and central target compared to other pathological markers.^3–5^ Mislocalisation of pathological tau to synaptic terminals is suggested to be an early pathological process that alters synaptic function and structure, eventually leading to synapse loss.^6–12^ Multiple models of acute^12^ and chronic tauopathy show synaptic dysfunction and/or degeneration, suggesting tauopathy is sufficient to induce synaptic changes.^13–15^

Excitatory neurotransmission is thought to be particularly vulnerable in tauopathy, evidenced by the loss of glutamatergic receptor expression in post-mortem brain tissue from patients.^16–18^ NMDA receptors (NMDAR) are a heavily implicated candidate in tauopathy pathogenesis, potentially via direct interactions between tau, NMDARs and Fyn kinase that cause NMDAR-mediated excitotoxicity.^19–22^ NMDAR subunit expression is also reduced in chronic models of tauopathy such as the rTg4510 mouse.^23^ Other alterations in glutamatergic receptors such as down and upregulation of AMPA receptor (AMPAR) subunit expression have also been observed in chronic tau overexpression models when characterised at different phases of the disease.^23–27^ Cultured neurons from the rTg4510 mouse model of tauopathy show tau overexpression leads to mislocalisation of AMPARs at synapses with an accompanying loss of synaptic function.^28^ This dysregulation could lead to deficits in synaptic transmission and synaptic plasticity, that have been observed in the rTg4510 model of tauopathy at a neurodegenerative phase of the disease.^29^ Altogether, chronic tauopathy mouse models show variable alterations in glutamatergic synaptic function, with increased, decreased and unchanged excitatory neurotransmission observed.^24–27,29^ Therefore, it is likely that tauopathy leads to physiological disruption at synapses, but there is a lack of evidence that links expression of synaptic proteins and receptors to function and how these may lead to differing effects with disease progression.

In addition to synapse loss, there is evidence that changes at the level of the dendrite occur to compensate for the loss of synaptic inputs.^30,31^ In advanced Alzheimer’s disease, changes in dendritic structure, and in particular dendritic atrophy, have been observed in post-mortem brain tissue.^32,33^ In models of tauopathy, there are varying dendritic phenotypes; both increased or decreased dendritic complexity are seen, likely reflecting changes across the progression of tauopathy and differences in tau isoform expression between models.^26,34,35^ These changes in dendritic structure may occur prior to the onset of frank neurodegeneration, perhaps to compensate for the loss of synapses and inputs to specific dendrites.^34,35^ Changes in dendritic structure could alter signal processing and synaptic integration within neurons, in addition to causing direct loss of input from lost synapses. Alternatively, functional preservation of specific afferent pathways has been reported in the rTg4510 mouse model of tauopathy, suggesting that pathology-driven changes in dendritic structure do not necessarily disrupt synaptic function.^27,36,37^ Rather, changes in dendritic complexity may preserve function or lead to compensatory alterations in tandem to changes in synaptic function at different disease phases.

In this study, we used the rTg4510 mouse model of tauopathy to explore how tauopathy can alter the receptor content and function of cortical synapses in early and progressed tauopathy. Abnormal tau inclusions are detectable in rTg4510 mice from ∼2 months of age, with visible atrophy in the cortex from ∼6 months of age.^38–41^ In addition to neurodegeneration, this model also exhibits large reductions in cortical synapse density and glutamatergic receptor expression by ∼8.5 months, by which time frank neurodegeneration is evident.^23,38–41^ Interestingly, prior to these dramatic changes in synaptic structures, aberrant dendritic spine stability was observed from 5-6 months of age in the somatosensory cortex, implying synaptic dysfunction occurs prior to synaptic degeneration in rTg4510 _mice.42,43_

Therefore, experiments were performed at 5.5 months and 7.5 months of age; rTg4510 mice at this later age point express marked cortical neurodegeneration.^38–41^ Using the somatosensory cortex as a neocortical model system, we isolated crude synaptosomes to measure synaptic marker peptide and glutamatergic receptor expression. Synaptic function was characterised using *in vitro* and *in vivo* whole-cell patch-clamp electrophysiological recordings, with contralateral whisker stimulation used to drive synaptic responses *in vivo*. Dendritic structure was examined from cells filled with biocytin during electrophysiological recordings.

## Materials and methods

### Ethical approval

Procedures were performed in accordance with the UK Animal (Scientific Procedures) Act 1986 and European Directive 2010/63/EU. Synaptosome extractions were performed at Eli Lilly and Co. and were approved by the Lilly UK Institutional Animal Care and Use Committee. Electrophysiological studies were performed at the Universities of Exeter and Bristol and were approved by their respective institutional Animal Welfare Ethical Review Bodies.

### Animals

Experiments used male rTg4510 and littermate control mice of 5.5 months (5.5 M; a total of WT: 35 mice, mean age 23 weeks; TG: 36 mice, mean age 22.6 weeks) and 7.5 M (a total of WT: 15 mice, mean age 33.6 weeks; TG: 16 mice, mean age 33 weeks).^38–40,44^ Mice were bred at ENVIGO (Oxon, UK) and kindly provided by Eli Lilly & Company. Mice were housed under standardised conditions (22 +/- 2 °C and 45 +/- 15 % humidity) on a 12-hour light/dark cycle with *ad libitum* access to food and water. For synaptosome analysis, brain homogenate from a P301S tauopathy mouse^45^ was used as a positive control. Experiments and analyses were performed blind to animal genotype.

### Synaptosome isolation

Mice were sacrificed by cervical dislocation. Brains were removed and the somatosensory cortex (+0.38 mm to −1.94 mm anteroposterior from Bregma) and cerebellum were isolated. Tissues were snap frozen and stored at −80 °C. Defrosted samples were sonicated for 10 sec (QSonica CL-18; 1 sec alternating on/off pulses, 40% pulse amplitude) in ice-cold buffer containing: 320 mM Sucrose, 10 mM EDTA, 50 mM Tris pH 7.4, protease inhibitor (cOmplete mini, Roche), Phosphatase Inhibitor Cocktail 2 (Sigma-Aldrich), Phosphatase Inhibitor Cocktail 3 (Sigma-Aldrich). Samples were centrifuged (Sigma 4-16KS, 12130 rotor) at 1,500 x g for 20 min at 4 °C. The supernatant was transferred to a new tube and centrifuged for 30 min at 16,000 x g at 4 °C. The supernatant was removed, and the pellet resuspended in 50 mM Tris containing phosphatase and protease inhibitors (TPP). Protein concentration was measured using a BCA protein assay and samples were normalised to 1 mg/ml in TPP.

### Protein electrophoresis

Samples were diluted in Sample Buffer (2x Laemmli, BioRad, or 4x NuPage LDS, Novex) containing 5% 2-mercaptoethanol, incubated in a heating block (Grant QBD2; 95 °C, 5 minutes), loaded into a NuPage 8% Bis-Tris Midi Gel (Invitrogen) in 1x MOPS SDS Running Buffer (Novex), and run at 150 V for 75 min (Invitrogen PowerEase 300W). Samples from different mouse ages and genotypes were distributed across gels in a randomised block design to minimise clustered variation. SeeBlue Plus 2 Prestained Standard (Invitrogen) was used as a molecular weight ladder.

Proteins were transferred by electroblotting (25 V, 80 min) onto a nitrocellulose membrane (0.45 µm pore size, Amersham Protran) soaked in NuPage transfer buffer (Novex) containing 20% methanol. Protein transfer was confirmed using Ponceau S (Sigma-Aldrich), and blots were cut into two at relevant molecular weights as needed. Membranes were blocked in PBS containing 0.05% Tween (PBS-T) and 5% dried milk (PBS-T-M), and incubated (overnight, 4 °C) with the primary antibody of interest (in PBS-T-M; see Table S1). Membranes were then washed (3 x 10 min in PBS-T) and incubated (1 hr) at room temperature (RT, ∼20°C) with corresponding secondary antibodies (in PBS-T-M; Table S1). Membranes were then washed (3 x 10 min in PBS-T). Proteins were incubated (5 min) with Dura or Femto substrates (ThermoScientific) and imaged using an Amersham Imager 600. If required, membranes were stripped using a stripping buffer (ThermoScientific), then re-incubated from the blocking stage.

### Synaptic protein analysis

Images were analysed using ImageQuantTL software (GE). Bands of interest were selected, and background signal subtracted using a rolling ball filter. Raw protein expression values were normalised within-lane to a housekeeper protein (GAPDH). Samples were excluded if the detected band (protein or housekeeper protein) was incomplete or visibly unclear.

### *In vitro* electrophysiology: recording

Mice were sacrificed by cervical dislocation at approximately the same time of day, brains were removed and placed into ice cold sucrose solution containing (in mM): 189 Sucrose, 10 D-Glucose, 26 NaHCO_3_, 3 KCl, 5 MgSO_4_.7H_2_O, 0.1 CaCl_2_, 1.25 NaH_2_PO_4_, bubbled in carbogen (95% O_2_, 5% CO_2_). 300 µm thick thalamocortical slices were prepared as described by Agmon & Connors, 1991 using a vibratome (Leica VT1200). Slices were stored at RT in artificial cerebrospinal fluid (aCSF) containing (in mM): 124 NaCl, 3 KCl, 24 NaHCO_3_, 2 CaCl_2_, 1.25 NaH_2_PO_4_, 1 MgSO_4_, 10 D-Glucose, bubbled in carbogen.

For recordings, slices were superfused with oxygenated aCSF (∼2 ml/min, ∼33°C) containing Gabazine (5 µM; Abcam). A bipolar stimulating electrode was placed in the middle of visible barrel fields in layer 4. Layer 2/3 putative pyramidal neurons in the same cortical column as the stimulating electrode were recorded using whole-cell patch clamp. Recording electrodes (4-7 MΩ) were filled with intracellular solution containing (in mM): 120 CsMeSO_4_, 6 NaCl, 5 QX314-Cl, 10 HEPES free acid, 10 BAPTA, 0.3, GTP.2Na salt, 4 ATP.Mg salt, 13.4 biocytin. Signals were amplified using an Axopatch 200B or Multiclamp 700A amplifier (Molecular Devices), digitised using an Axon Digidata 1550 (Molecular Devices) and recorded using pClamp v10.4 software (Molecular Devices). Signals were lowpass filtered at 5 or 10 kHz and sampled at 20 kHz.

Liquid junction potential error (-15 mV) was corrected for. Series resistance was measured at the start of recordings. Evoked excitatory postsynaptic currents (EPSCs) were recorded in voltage-clamp following electrical stimulation of L4 (1 ms duration; intensity adjusted to generate 50-150 pA EPSCs). AMPAR and NMDAR currents were measured at holding potentials of −70 mV and +40 mV, respectively. L689560 (NMDAR antagonist, 5 µm; Abcam) was used to confirm the NMDAR-dependency of +40 mV responses in a subset of experiments. EPSC paired pulse ratio (PPR) was measured in voltage-clamp at −70 mV in response to six stimuli at 5 Hz, 10 Hz, or 30 Hz, delivered every 15 seconds.

### *In vitro* electrophysiology: analysis

Analyses were performed using MATLAB or Clampfit v10.4 software. Cells with series resistance >40 MΩ (>95th percentile of the distribution) were excluded. Series resistance was lower in the 7.5 M age group (WT mean: 17 +/- 0.6 MΩ, TG mean: 16 +/- 0.7 MΩ) compared to 5.5 M (WT mean: 22 +/- 0.7 MΩ, TG mean: 22 +/- 0.8 MΩ; genotype effect: p=0.97, age effect: χ2_(1,227)_=17.4, p<0.005, interaction: p=0.37). Therefore, datasets were compared within but not between age group. AMPAR responses were measured as the peak response at −70 mV. NMDAR responses were measured 50 ms post stimulation at +40 mV. NMDA:AMPA ratio was calculated by dividing NMDAR responses by AMPAR responses. PPR was calculated by dividing the peak response of the second evoked EPSC by the peak response of the first evoked EPSC.

### Histology

After recordings, slices were fixed (overnight, 4°C) in 4% paraformaldehyde in 0.1M PBS, transferred to 0.1M phosphate buffer (PB) and stored at 4°C. Antigen retrieval was performed by incubating slices in dH2O at 78°C for 25 min. After cooling (35 min), slices were washed in 0.1M PB (3 x 5 min) and 0.1M PBS (3 x 5 min). Slices were first blocked (30 min, RT) in 100 mM glycine in 0.1M PBS containing 0.3% Triton-X and 0.05% sodium azide (PBS-T-A), and then blocked (1 hr, RT) in 0.5% normal goat serum (Vector Labs) in PBS-T-A. Slices were then incubated (3 days, 4°C) in MC-1 primary antibody^47^ (1:1,000 in PBS-T-A). Slices were washed (3 x 5 min) in 0.1M PBS and incubated (overnight, 4°C) with goat anti-mouse Alexa Fluor 594 IgG secondary antibody and anti-avidin Alexa Fluor 488 conjugate (both 1:1,000 in PBS-T-A; ThermoFisher). Slices were washed (4 x 30 min) in 0.1M PBS, rinsed (5 min) in dH2O, and mounted using VectaShield (Vector Labs). Slices were imaged using two-photon (Scientifica) or confocal (Lecia SP8) microscopy. Z-stacks (0.5 µm optical slice) were taken of neuronal structure and analysed using Simple Neurite Tracer and Sholl analysis (5 µm bins) plugins in Fiji software.^48,49^

### *In vivo* electrophysiology: recordings

Experimental surgeries were all performed at a similar time of day. Mice were anaesthetised using isoflurane (induction: 3-4%, maintenance: 1-2%) and fixed in a custom stereotaxic frame with removable baseplate. Body temperature was maintained at 37 °C using a homeothermic blanket. An incision was made in the scalp and a head-fixation bar was affixed to the occipital skull plate. A dental cement well was made around somatosensory cortex and filled with aCSF containing (in mM): 135 NaCl, 5 KCl, 5 HEPES free acid, 1.8 CaCl_2_, 1 MgCl_2_. A small craniotomy was drilled at coordinates (from Bregma): +1.3 mm anteroposterior, −3.4 mm mediolateral, and the skull flap and dura were removed under aCSF. Mice received a subcutaneous dose of urethane (1.5-2 mg/kg) to maintain anaesthesia, and isoflurane was slowly withdrawn.

Mice were transferred via the baseplate to a two-photon microscope (Scientifica) equipped with Ti:Sapphire laser (MaiTai, Spectra-Physics). The whiskers contralateral to the recording hemisphere were bundled using Vaseline and a custom stimulator (1 mm aperture attached to a pneumatic picopump; World Precision Instruments) was positioned ∼5 mm from the whisker bundle. The brain surface was visualised under white-light, and a glass recording electrode (4-10 MΩ) was inserted into the brain. The electrode internal solution contained (in mM): 135 mM K-Gluconate, 5 mM NaCl, 10 mM HEPES free acid, 0.2 mM EGTA, 0.3 mM GTP-Na salt, 4 mM ATP-Mg salt, 6.7 mM Biocytin, 25 μM Alexa Fluor 594 Hydrazide. Two-photon imaging was used to target patch-clamp recordings to putative layer 2/3 (L2/3) pyramidal neurons using the shadow patch method.^50,51^ Some recordings were performed with MK801 (NMDAR antagonist, 1 mM; Hello Bio) in the intracellular solution. Signals were amplified using a Multiclamp 700A, digitised using a Power 1401 (CED), and recorded using Signal v5.07 software (CED). Raw signals were lowpass filtered at 10 kHz and sampled at 50 kHz. Supplementary doses of urethane were given to maintain a stable anaesthetic plane throughout recordings.

Recordings were not corrected for liquid junction potential error and bridge balance was not applied. After achieving whole-cell configuration, the series resistance was measured in voltage-clamp. Then, 3 minutes gap-free recording was performed in current-clamp (I=0). Air-puffs (10 mBar, 100 ms duration) were delivered to stimulate the whiskers (5 Hz, 1 s) in the rostral-caudal direction, every 30 seconds for 15 minutes. Vm was recorded throughout stimulation in current-clamp (I=0). Series resistance was re-checked before and after whisker stimulation.

### *In vivo* electrophysiology: analysis

Neurons were excluded if membrane voltage (Vm) exceeded 0 mV for >1 second, if series resistance >100 MΩ, or Vm was recorded <6 minutes during whisker stimulation. To isolate synaptic responses to whisker stimulation, action potentials (AP) were removed by normalising values of Vm exceeding AP threshold (dVm/dt >30 mV/ms) to the AP threshold. ^52,53^ Responses were split into those occurring during Up or Down states. Down states were defined as epochs of hyperpolarisation with Vm range <3 mV in the 50 ms preceding whisker stimulation.^54^ Signals not meeting these criteria were classified as Up states.^54^ Up and Down state responses were normalised to the mean membrane potential (Vm) in first 5 ms post stimulation.^54,55^ Synaptic responses to the first whisker stimulus and 5 Hz train were analysed. For first stimuli, the peak response was the max Vm within 0-40 ms post stimulus onset; the secondary postsynaptic potential (PSP) component was the mean Vm between 50-100 ms post stimulus onset. For the 5 Hz train, the area under the curve (AUC) from stimulus onset (0 ms) to 200 ms after the last stimulus was calculated to measure the total response envelope. The cross-correlation was calculated between the stimulus train and Z-scored Vm during Down states.

### Statistical Analysis

Datasets were analysed using generalised linear mixed models (GLMMs) using R (v3.5.1) with RStudio (v1.1.456). Data were visually inspected to identify appropriate distributions, link functions and factors. GLMMs included random factors to control for statistical bias and pseudo-replication.^56–58^ The intraclass coefficient (ICC) evaluated whether random factors (e.g. animal, gel blot) significantly affected variance. Where ICC<0.1 (0-1 scale), random factors were removed to avoid overfitting. The data dispersion, homogeneity of variance, Akaike information criterion, and normality of residuals were used to evaluate GLMM fit. Statistical significance (p<0.05) of factors in the GLMM was determined using goodness of fit chi squared (χ2) tests. Post hoc tests were performed using Tukey-Kramer multiple comparisons adjustment. Descriptive statistics are reported as mean ± SEM. Data are typically presented as box plots: squares denote the mean; boxes define ± standard error of the mean (SEM); horizontal lines denote the median; dots plot individual samples. Line graphs plot mean ± SEM.

### Data availability

The data and MATLAB scripts used for this study are available from the corresponding author, upon reasonable request.

## Results

### Reduced synaptic protein expression in the somatosensory cortex of rTg4510 mice at 5.5 M and 7.5 M of age

To assess the effect of pathological tau on levels of synaptic proteins, crude synaptosomes were isolated from the somatosensory cortex of TG and WT littermate mice at two different ages: 5.5 M (10 WT, 11 TG) and 7.5 M (10 WT, 11 TG). Protein expression levels in synaptosomes were quantified by Western blot. The presynaptic protein synaptophysin was significantly reduced in TG synaptosomes compared to WT synaptosomes (χ^2^_(1,42)_=4.73, p=0.02*, ICC=0.53), with no age effect (p=0.64) or interaction between genotype and age (p=0.58) (*Figure 1A&B*). The excitatory postsynaptic scaffold protein, PSD95, was also significantly reduced in TG synaptosomes (genotype: χ^2^_(1,40)_=14.13, p<0.005***, ICC=0.39), with a significant age effect (age: χ^2^_(1,40)_=3.75, p=0.05*, ICC=0.39; *Figure 1A&C*). There was no interaction between genotype and age (p=0.31). The reduced expression of synaptophysin and PSD95 in TG samples suggests synapse loss and/or weakening is an early feature of tauopathy, with more substantial effects on postsynaptic structures.

**Figure 1.**
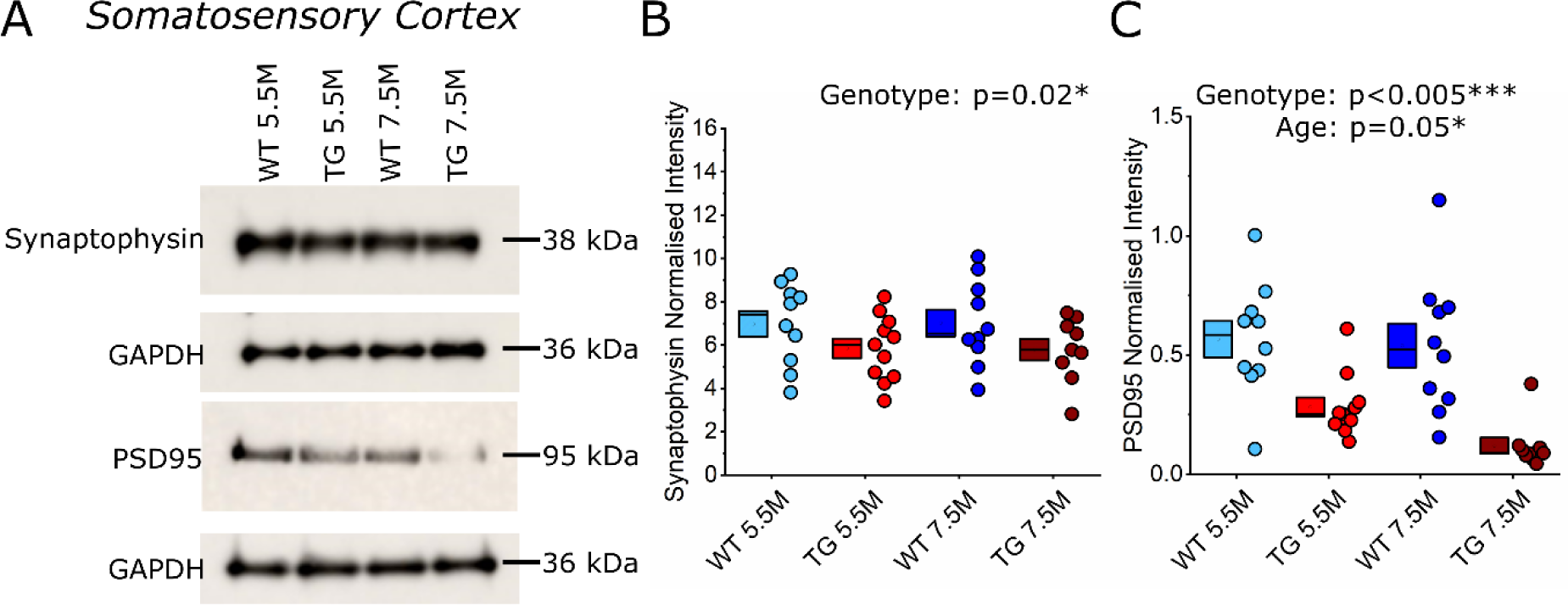
Expression of the synaptic markers synaptophysin and PSD95 were reduced in the somatosensory cortex in 5.5 M and 7.5 M rTg4510 mice. **A)** Representative Western blots of synaptophysin and PSD95 expression in the somatosensory cortex from WT and TG animals at 5.5 and 7.5 M (labelled along the top). Annotated along the left-hand side is the detected protein marker for the blot shown in the middle. Annotated on the right-hand side is the corresponding molecular weight of the protein. Expression levels for each protein were normalised to the corresponding GAPDH band (below). **B&C)** Box and dot plots showing quantification of each marker. Statistics: GLMM, main significant effects labelled on the graph, with statistical significance denoted by asterisks (p<0.05*, p<0.005***). **B)** Quantification of synaptophysin protein expression, normalised to GAPDH. **C)** Quantification of PSD95 protein expression, normalised to GAPDH.

### Glutamatergic receptor expression was reduced in rTg4510 mice at 5.5 M and 7.5 M of age

The expression of the AMPAR subunits GluA1, GluA2, and GluA3 were measured, as these are the most prevalent AMPAR subunits in the adult rodent brain.^59–61^ There was no significant effect of genotype (p=0.26), age (p=0.11) or interaction between genotype and age (p=0.44) on GluA1 expression (*Figure 2A&B*). In contrast, there was a substantial reduction in GluA2 (χ^2^_(1,42)_=29.1, p<0.005***, ICC=0.06; *Figure 2A&C*) and GluA3 expression (χ^2^_(1,41)_=9, p<0.005***, ICC=0.25; *Figure 2A&D*) within TG synaptosomes compared to WT. This result mirrors the loss of PSD95 observed in TG mice; fewer postsynapses or less postsynaptic scaffolding may lead to fewer postsynaptic AMPARs. There was also a significant age dependent decrease of GluA2 expression (χ^2^_(1,42)_=4.33, p=0.04*, ICC=0.06), which was not seen for GluA3 (p=0.15). There was no interaction between genotype and age for either GluA2 (p=0.43) or GluA3 (p=0.5) expression. GluN1 subunit expression was also quantified to assess NMDAR expression within synaptosomes. There was a significant effect of genotype (χ^2^_(1,42)_=6.27, p=0.01**, ICC 0.27) and age (χ^2^_(1,42)_=5.72, p=0.02*, ICC=0.27) on the expression of GluN1, with no interaction between the two factors (p=0.62) (*Figure 2A&E*). Overall, this suggests that loss of the synaptic proteins synaptophysin and PSD95 is complemented by decreased AMPAR and NMDAR expression in TG mice at both 5.5 M and 7.5 M ages.

**Figure 2.**
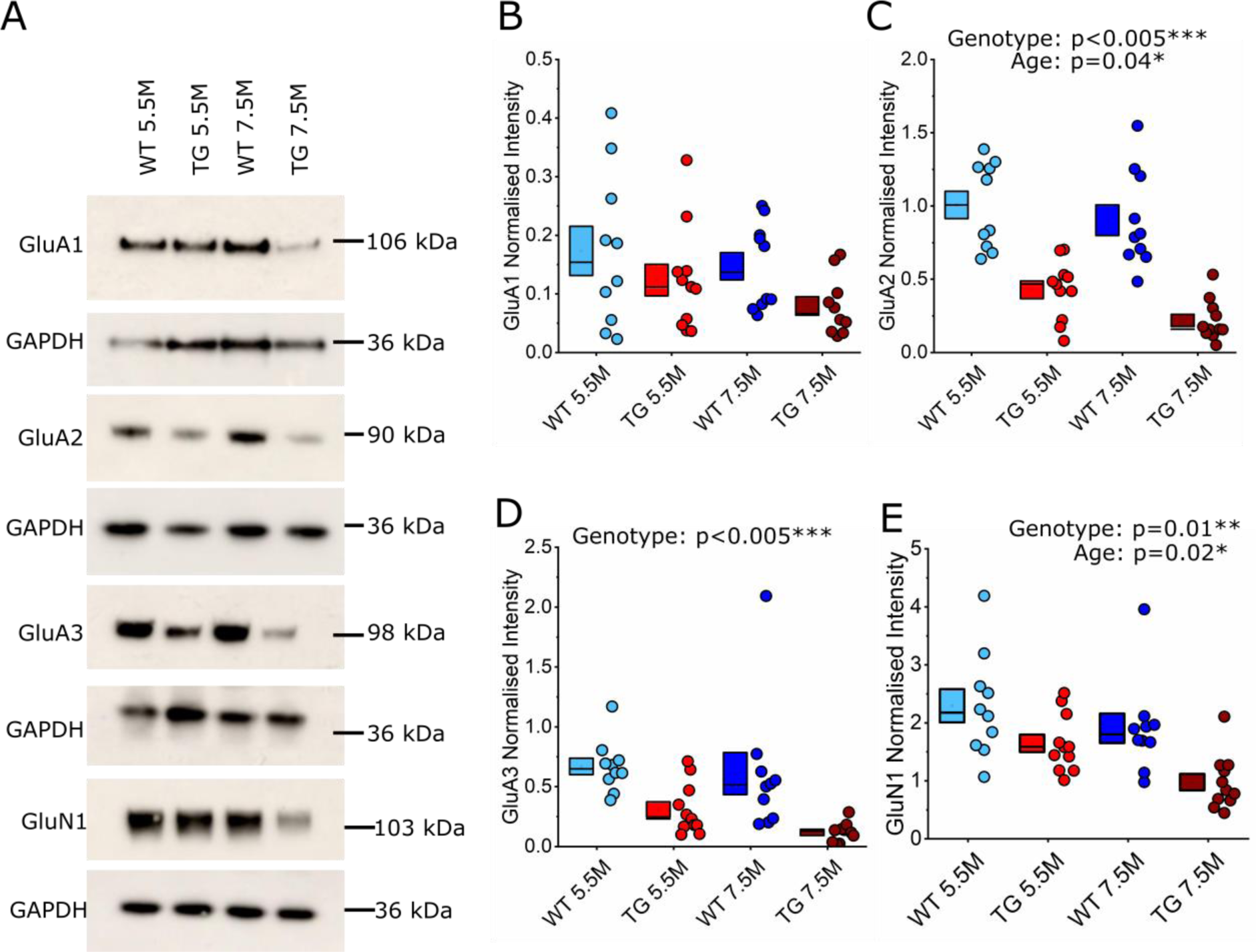
AMPAR (GluA2 and GluA3) and NMDAR (GluN1) subunit expression was reduced in the somatosensory cortex in 5.5 M and 7.5 M rTg4510 mice. **A)** Representative Western blots of GluA1-3 and GluN1 expression in the somatosensory cortex of WT and TG mice at 5.5 and 7.5 M. Annotated on the left-hand side is the detected protein marker for the blot shown in the middle. Annotated on the right-hand side is the corresponding molecular weight of the protein. Expression levels for each protein were normalised to the corresponding GAPDH band (below). **B-E)** Box and dot plots showing quantification of each marker. Statistics: GLMM, main significant effects labelled on the graph, with statistical significance denoted by asterisks (p<0.05*, p<0.005***). **B)** Quantification of GluA1 expression, normalised to GAPDH. **C)** Quantification of GluA2, normalised to GAPDH. **D)** Quantification of GluA3, normalised to GAPDH. **E)** Quantification of GluN1 expression, normalised to GAPDH.

### *In vitro* assessment of synaptic function: paired pulse ratio was unaltered in rTg4510 mice

To assess cortical synaptic function, we performed whole-cell patch clamp recordings of L2/3 pyramidal neurons in somatosensory cortex brain slices from TG and WT littermate mice at 5.5 M & 7.5 M. Short-term synaptic plasticity, which depends on presynaptic release probability, was assessed by measuring the PPR in response to electrical stimulation of L4-to-L2/3 synapses. PPRs of evoked EPSCs were measured in response to 5 Hz, 10 Hz, and 30 Hz stimulation. These synaptic activation frequencies span theta and gamma band ranges, which are associated with the temporal organisation of synaptic innervation *in vivo*.^62–66^

For the 5.5 M age group (29 WT neurons, 8 animals; 27 TG neurons, 9 animals), lower stimulation frequencies (5 Hz and 10 Hz) produced a paired pulse depression of EPSC amplitude in both WT and TG neurons (PPR of 0.85 ± 0.05 in WT cells and 0.9 ± 0.09 in TG cells at 5 Hz, genotype effect p=0.36, *Figure 3A*; 0.94 ± 0.08 in WT cells and 0.92 ± 0.09 in TG cells at 10 Hz, genotype effect p=0.84, *Figure 3B*). At 30 Hz, paired stimuli drove a facilitating PPR that was similar in both WT and TG neurons (PPR of 1.26 ± 0.17 in WT cells and 1.28 ± 0.26 in TG cells, genotype effect p=0.88, *Figure 3C*). There was no significant effect of genotype on any short-term plasticity measures at 5.5 M.

**Figure 3.**
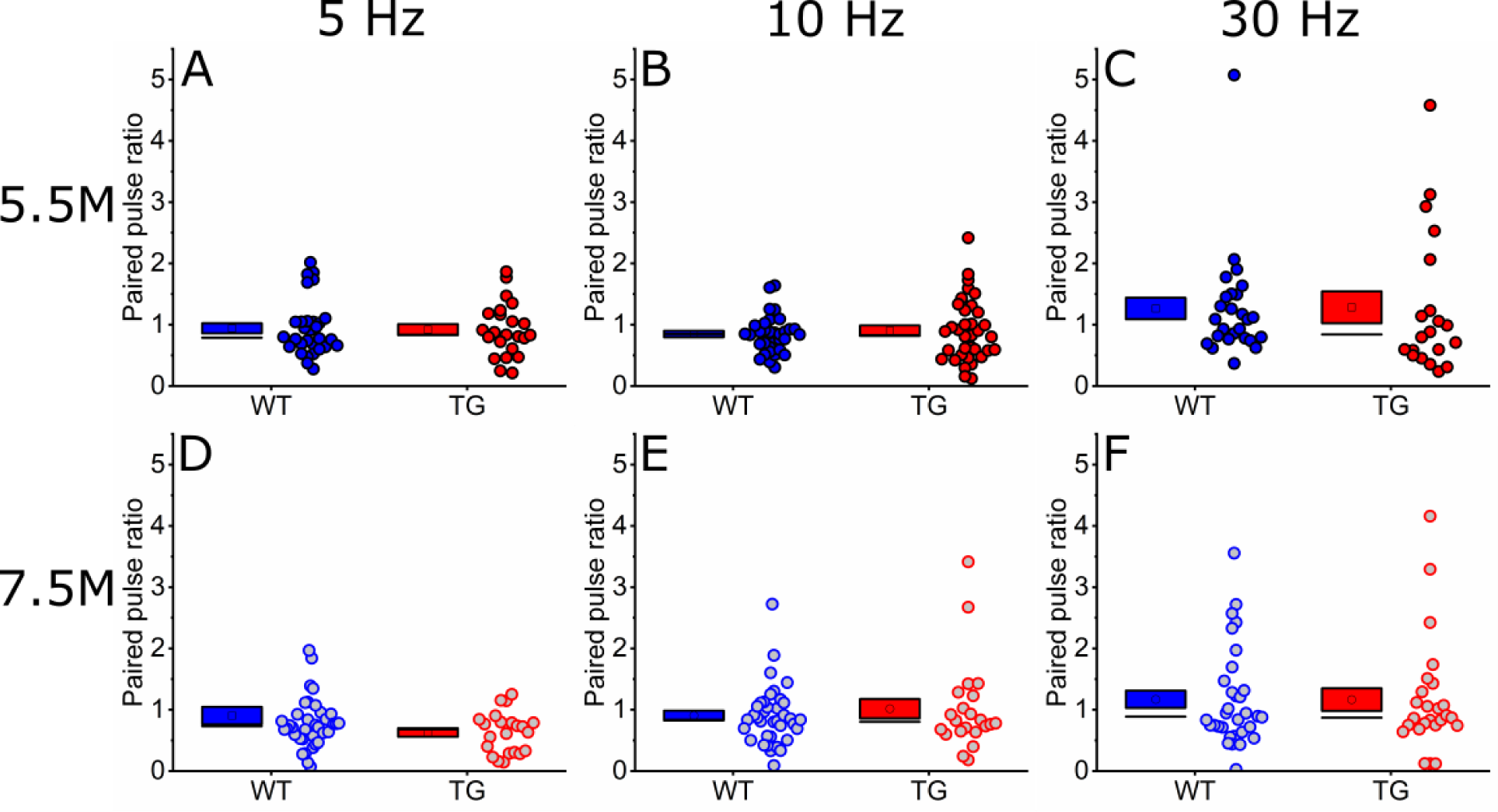
Paired pulse ratios were similar between 5.5 M and 7.5 M WT and TG mice. PPRs for both 5.5 M and 7.5 M age groups (5.5 M, **A-C**; 7.5 M, **D-F**) for all three tested stimulation frequencies (5 Hz, **A&D**; 10, Hz **B&E**; 30 Hz, **C&F**). PPR values above 1 suggest an overall facilitation of the response; PPR values below 1 suggest a depressing PPR. Note that the mean PPR is typically below 1 for synaptic responses to 5 Hz and 10 Hz stimulation, and above 1 for synaptic responses to 30 Hz stimulation. Statistics: GLMM, no significant differences observed.

In 7.5 M animals (38 WT neurons, 19 TG neurons, 5 animals/genotype), lower stimulation frequencies (5 Hz and 10 Hz) produced similar responses to 5.5 M. 5 Hz stimulation produced a depressing PPR, whilst 10 Hz stimulation produced a depressing PPR for the WT group and a small facilitating PPR for the TG group (mean PPR of 0.9 ± 0.14 in WT cells and 0.63 ± 0.08 in TG cells at 5 Hz, genotype effect p=0.21, *Figure 3D*; mean PPR of 0.9 ± 0.09 in WT cells and 1.05 ± 0.17 in TG cells at 10 Hz, genotype effect p=0.89, Figure 3E). At 30 Hz, the mean PPR was facilitating in both WT and TG neurons (mean PPR of 1.16 ± 0.16 in WT cells and 1.26 ± 0.2 in TG cells at 30 Hz, genotype effect p=0.95, *Figure 3F*). Overall, there was no significant effect of genotype on short-term synaptic plasticity measures at either age. This suggests that tauopathy does not affect neurotransmitter release probability at L4-to-L2/3 synapses, suggesting that presynaptic release is unaltered by tauopathy at this disease phase.

### *In vitro* assessment of synaptic function: NMDA:AMPA receptor response ratio was unaltered in rTg4510 mice

Decreased AMPAR and NMDAR subunit expression (*Figure 2*) could cause changes in the synaptic responses mediated by these receptors. Therefore, we tested the functional receptor content of L4-to-L2/3 synapses by measuring the NMDA:AMPA receptor EPSC ratio. There was no significant effect of genotype at 5.5 M (χ2_(1,63)_=0.1, p=0.75; *Figure 4A&B*; n=39 WT cells, n=33 TG cells, 10 animals/genotype) or at 7.5 M (χ2_(1,72)_=0.12, p=0.74; *Figure 4C&D*; n=42 WT cells, n=30 TG cells, 5 animals/genotype). This suggests that L4-to-L2/3 synapses maintain relative levels of functional postsynaptic glutamate receptors despite putative synapse loss during progressive tauopathy.

**Figure 4.**
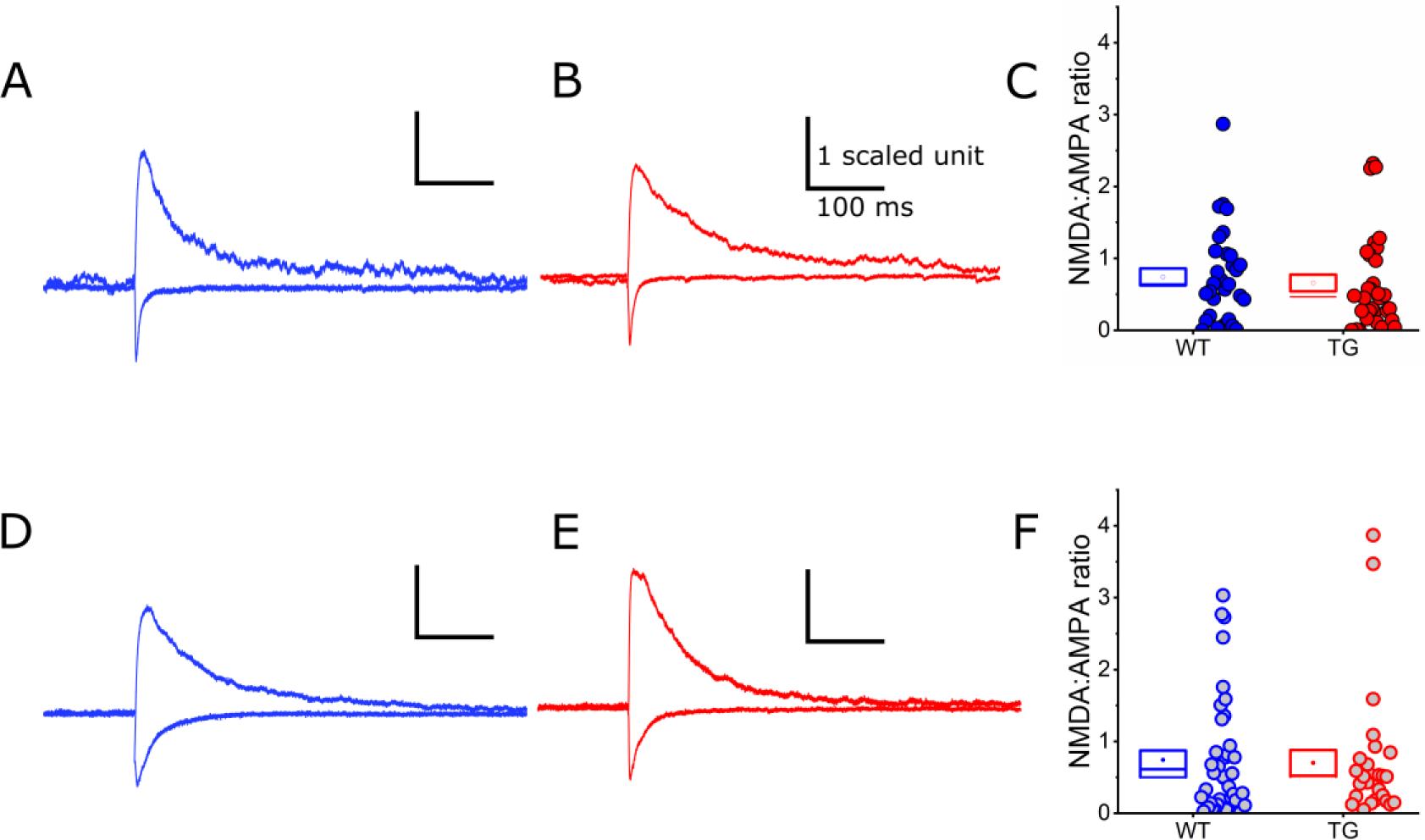
The NMDA:AMPA receptor response ratio was similar between 5.5 M and 7.5 M WT and TG mice. **A, B, D, E)** Example waveforms of NMDA and AMPA receptor responses, normalised to the AMPA receptor peak component (WT blue **A&D**, TG red **B&E**). The stimulus artefact has been removed from each waveform. **A&B)** Waveforms from WT and TG animals at 5.5 M of age. **C)** Box and dot plots showing pooled NMDA:AMPA receptor ratios at 5.5 M. **D&E)** Waveforms from WT and TG animals at 7.5 M of age. **F)** Box and dot plots showing pooled NMDA:AMPA receptor ratios at 7.5 M. Statistics: GLMM, no significant differences observed.

### Cortical pyramidal neurons exhibited increased dendritic branching proximal to the soma in 5.5 M rTg4510 mice

Since decreased expression of synaptic proteins in somatosensory cortex (*Figure 1 & Figure* 2) was not associated with changes in glutamatergic synaptic transmission in 5.5 M TG mice *(Figure 3 & Figure* 4), we assessed whether changes in neuronal structure may compensate for altered synapse density and/or receptor subunit composition.^27^ Sholl analysis was used to compare dendritic reconstructions of neurons from the 5.5 M age group that were labelled with biocytin during electrophysiological recordings (21 WT neurons, 10 animals; 26 TG neurons, 13 animals; *Figure 5A&B*).

**Figure 5.**
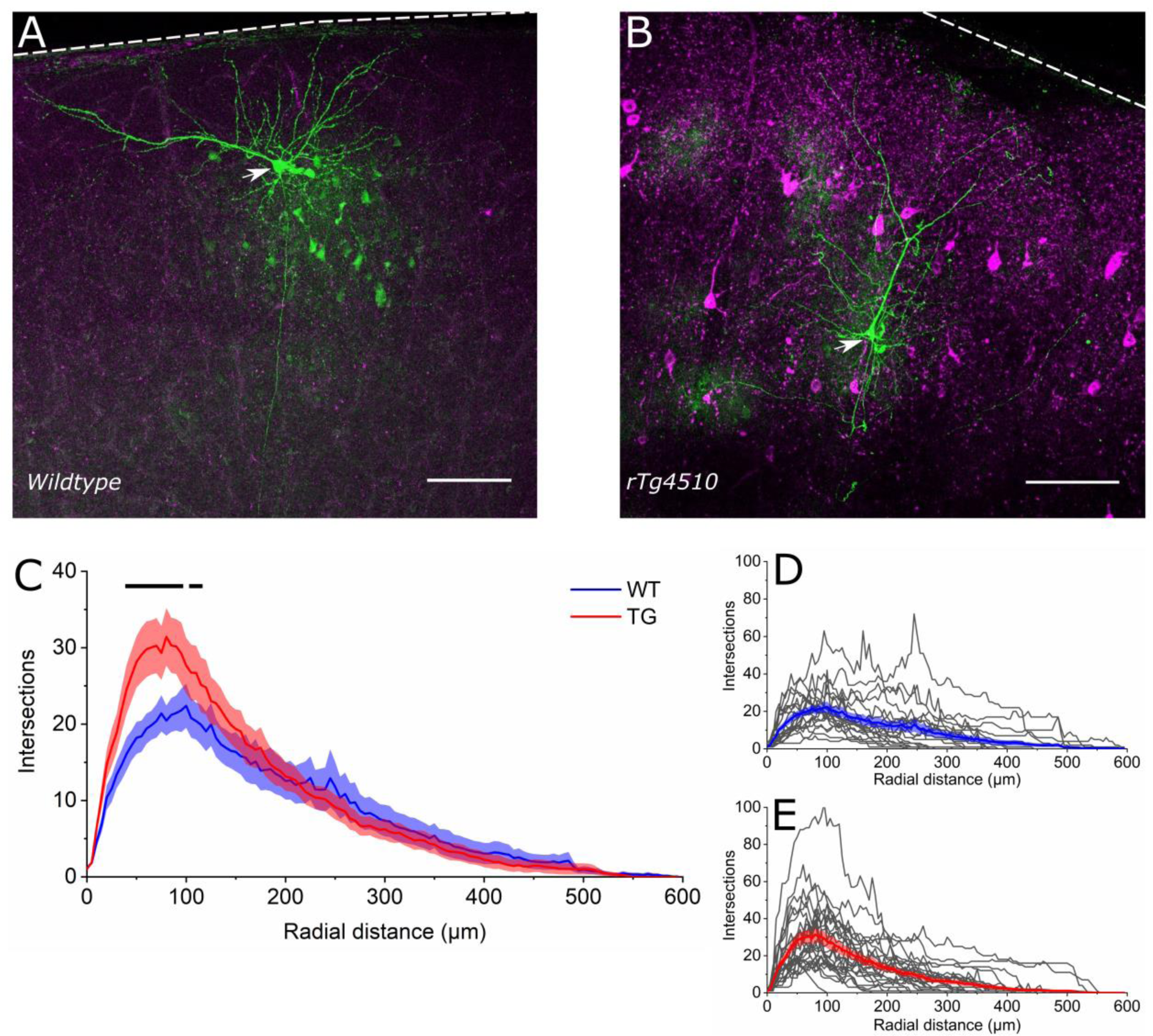
Restructuring of proximal dendrites in rTg4510 cortical pyramidal neurons at 5.5 M. **A&B)** Photomicrographs of representative morphologically recovered WT and TG neurons (green, Alexa Fluor 488). The soma of each recovered neuron is denoted by a white arrowhead. Immunolabelling for conformationally altered tau (MC-1 antibody) is shown in magenta. Note that anti-tau immunolabelling is absent in WT tissue **(A)** but is prolific in TG tissue **(B)**. Scale bar: 100 μm. The dashed white line denotes the location of the pia. **C)** A Sholl plot of the average intersections per radial distance of WT (blue) and TG (red) cortical putative pyramidal neurons. Thick lines represent the mean, with shading representing the SEM. The black line above the plot denotes statistical significance at p<0.05. Accurate statistically significant p values (per radial distance, in μm) are: 35, p=0.02; 40, p=0.005; 45, p<0.005; 50, p<0.005; 55, p<0.005; 60, p<0.005; 65, p<0.005; 70; p<0.005; 75, p=0.008; 80, p<0.005; 85, p<0.005; 90, p<0.005 95, p=0.01; 105, p=0.05; 110, p=0.03. **D&E)** Sholl plots for all morphologically reconstructed WT **(D)** and TG neurons **(E)**. Thick lines represent the mean.

This analysis revealed a significant main effect of radius (χ^2^_(119,5520)_=5715, p<0.005***), but not of genotype (p=0.89). There was also a significant interaction between genotype and radius, suggesting that TG neurons have a different dendritic structure from WT neurons in specific regions of the dendrite (χ^2^_(119,5520)_=310.4, p<0.005***). Specifically, TG neurons had significantly more dendritic branch intersections in the region proximal to the soma (within 35 – 110 µm; *Figure 5C*). Therefore, whilst there was no overt dysfunction of L4-to-L2/3 synaptic transmission in 5.5M TG mice, there may be altered network-level integration of synaptic input across the dendritic tree due to dendritic restructuring.

### *In vivo* assessment of synaptic function: evoked synaptic responses had an increased peak amplitude in 5.5 M rTg4510 mice

L2/3 somatosensory cortex pyramidal neurons receive differentially distributed synaptic input onto the dendritic tree, with the distal apical tuft receiving long-range and higher order inputs from distal areas whilst the proximal dendrites receive local L4, L2/3, and first-order thalamic inputs.^67–70^ Therefore, alterations in dendritic structure may disrupt the integration of synaptic inputs with complex spatiotemporal distribution across the dendrite. To assess this, we used *in vivo* whole-cell patch clamp recordings of L2/3 barrel (i.e. whisker somatosensory) cortex pyramidal neurons in urethane anaesthetised mice to measure functional synaptic integration at 5.5 M (n=15 animals/group). Air puffs were used to generate rhythmic (5Hz for 1s) rostrocaudal deflections of the contralateral whiskers.

Under urethane anaesthesia, neuronal Vm cycles between Up and Down states, which impacts the response to incoming synaptic drive.^71–74^ Up-Down state oscillations were consistently observed in recordings from WT and TG neurons (*Figure 6A*), so stimulus-evoked PSPs were separated based on whether the onset of whisker deflection coincided with an Up or Down state. Whisker deflection generated complex PSP waveforms combining depolarising and hyperpolarising phases that suggested integration of multiple excitatory and inhibitory synaptic conductances (*Figure 6B-E*). First, the response to the initial whisker deflection was analysed by measuring the peak Vm immediately after whisker deflection and the average PSP between 50-100 ms post deflection, which corresponds to a period of integration of polysynaptic inputs triggered by the whisker deflection.^75^

**Figure 6.**
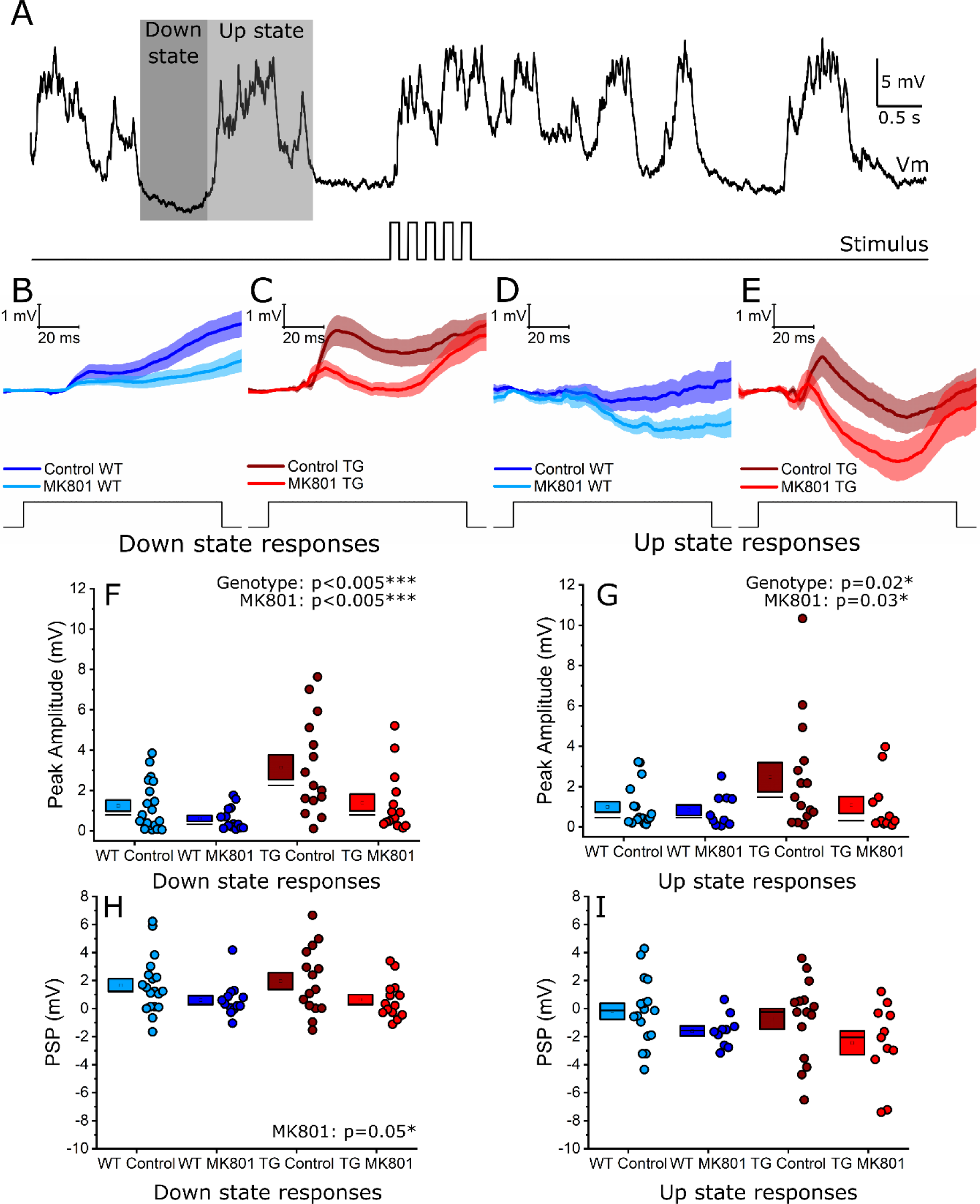
Sensory stimulation elicited a larger initial synaptic response in rTg4510 cortical pyramidal neurons at 5.5 M. **A)** An example voltage trace showing Up states (light shading) and Down states (dark shading) in an *in vivo* whole-cell recording of a putative pyramidal neuron in somatosensory cortex. An example synaptic response evoked by rhythmic (5 Hz, 1s) whisker deflection is also illustrated; the stimulus time course is illustrated beneath the voltage trace. **B&C)** Averaged synaptic responses to the first whisker deflection of the 5 Hz train during Down states in WT **(B)** and TG **(C)** neurons. Both control and MK801 conditions are shown. Thick lines represent the mean, with shading representing SEM. **D&E)** Averaged synaptic responses to the first whisker deflection of the 5 Hz train during Up states in WT **(D)** and TG **(E)** neurons. Both control and MK801 conditions are shown. Thick lines represent the mean, with shading representing SEM. **F&G)** Quantification of the initial (0-40 ms post stimulation) peak of the synaptic response evoked by whisker deflection during Down states **(F)** and Up states **(G)**. **H&I)** Quantification of the average PSP 50-100 ms after whisker deflection during Down states **(H)** and Up states **(I)**. Statistics: GLMM, main significant effects labelled on the graph, with statistical significance denoted by asterisks (p<0.05*, p<0.005***).

The peak of the whisker deflection-evoked response was significantly larger in TG neurons compared to WT controls during both Down states (genotype effect: χ^2^_(1,62)_=10.7, p<0.005***, ICC=0.11) and Up states (genotype effect: χ^2^_(1,53)_=5.2, p=0.02*, ICC=0.23; *Figure 6F&G*). Interestingly, intracellular application of the use-dependent NMDAR antagonist MK801 via the recording pipette significantly reduced the peak response during both Down and Up states (Down state MK801 effect: χ^2^_(1,62)_=10.4, p<0.005***; Up state MK801 effect: χ^2^_(1,53)_=4.7, p=0.03*; *Figure 6F&G*). This shows that postsynaptic NMDARs contributed to the initial whisker deflection-evoked depolarisation, but there was no difference in NMDAR contribution between WT and TG neurons in either Down (p=0.91) or Up (p=0.25) states.

The average PSP 50-100 ms post whisker-deflection was not different between genotypes during Down states (p=0.61) or Up states (p=0.53; *Figure 6H&I*). As expected, MK801 treatment affected PSP amplitude, significantly decreasing the PSP during Down states (MK801 effect: χ^2^_(1,62)_=3.8, p=0.05*; *Figure 6H*) with no effect on Up state responses (p=0.08; *^F^igure 6^I^*). There was no interaction between genotype and MK801 treatment on PSP responses during Down (p=0.82) or Up (p=0.79) states.

*In vivo* assessment of synaptic function: the compound synaptic charge evoked by rhythmic whisker stimulation was not altered in 5.5 M rTg4510 mice

Next, the response to 5 Hz whisker deflection was examined. This stimulus generated an envelope of synaptic activity that could be used to examine integration of sensory drive. Thus, the area under the curve (AUC) of the entire PSP was measured to assess overall synaptic activation. During Down states, there was no significant effect of genotype (p=0.85), or MK801 (p=0.29), and no interaction between genotype or MK801 treatment (p=0.37; *Figure 7E*). This was also the case for responses initiated during Up states, with no significant effect of genotype (p=0.98), MK801 treatment (p=0.24), or interaction between genotype or MK801 treatment (p=0.35; *Figure 7F*). Therefore, despite genotype- and MK801-dependent differences in the response to initial whisker deflection (*Figure 6*), the compound synaptic activation evoked by a stimulus train was similar between genotypes and with MK801 treatment.

**Figure 7.**
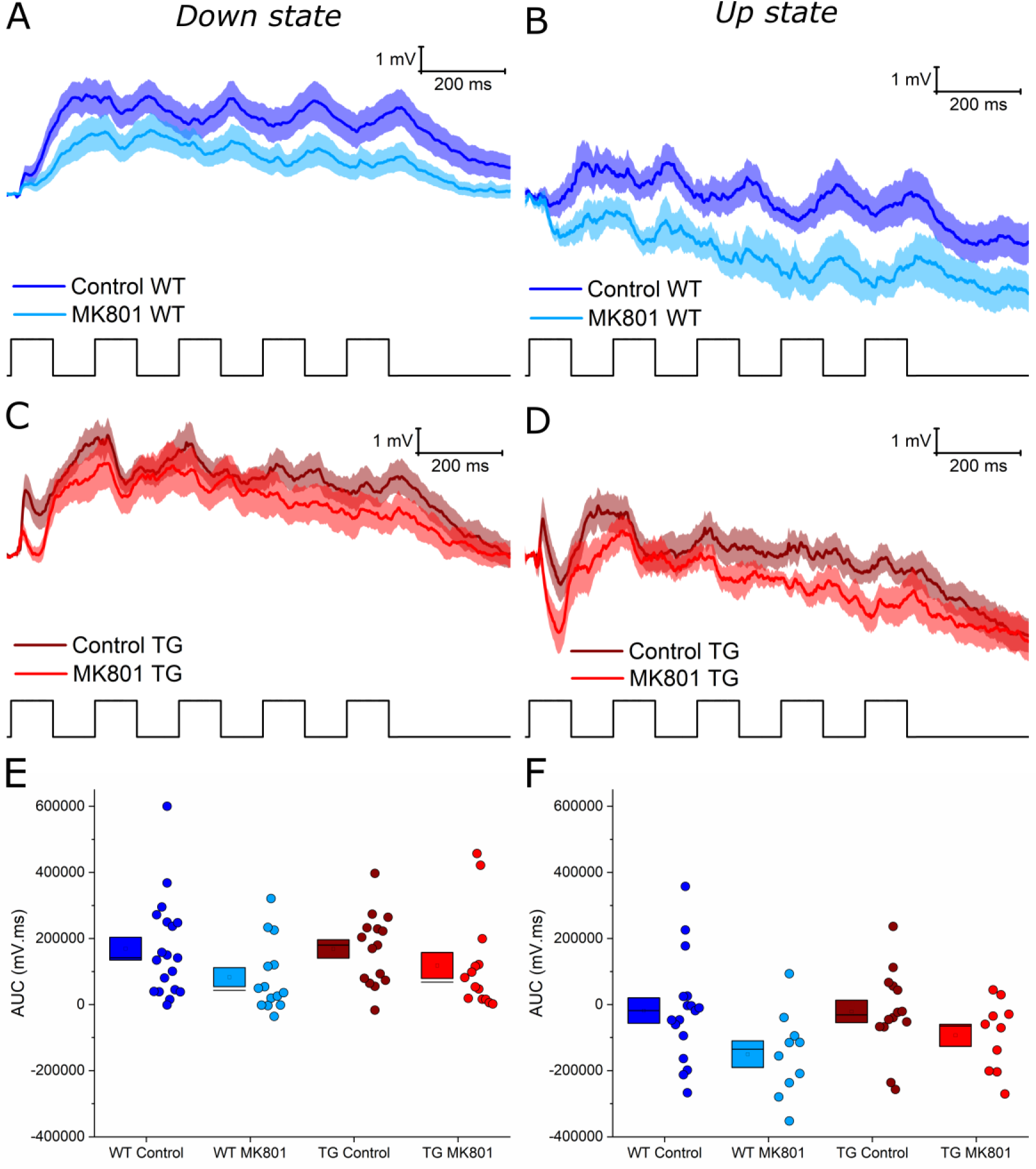
Genotype or MK801 treatment did not alter the compound synaptic charge evoked by 5 Hz sensory stimulation. **A-D)** Average compound synaptic responses evoked by the full 5 Hz train of whisker stimuli during Down states **(A&C)** and Up states **(B&D)** for WT **(A&B)** and TG **(C&D)** neurons. Thick lines represent the mean, with the shading representing the SEM. **E&F)** Pooled data illustrating the AUC of the stimulus train-evoked compound synaptic response during Down states **(E)** and Up states **(F)** states. Statistics: GLMM, no significant differences observed.

### Reduced temporal coordination of *in* vivo synaptic responses to rhythmic whisker stimulation in 5.5 M rTg4510 mice

Whilst compound synaptic activation was similar between genotypes, inspection of WT and TG responses to 5 Hz whisker deflection (*Figure 7A-D*) indicated that the temporal profile of the response was altered between genotypes. Cross-correlation analysis was performed between the 5 Hz stimulus signal and the synaptic response to determine how faithfully the response followed the input pattern *(Figure 8A&B).* Due to differences in driving force between Up and Down states leading to differences in the response size, and to align with previous literature on cortical synaptic responses to whisker stimulation-evoked activity, analysis was performed on Down state responses.^73,75–77^ This revealed clear loss of rhythmicity in TG synaptic responses compared to WT controls (*Figure 8E&F;* peak cross-correlation: χ^2^_(1,62)_=8.2, p<0.005; *Figure 8C-E*). Measurement of the peak correlation lags to assess if there was a shift in stimulus-response timing revealed no genotype effect (p=0.27; *Figure 8F*), suggesting the peak stimulus-response correlation occurred at a similar time in WT and TG neurons. There was no effect of MK801 on the peak correlation coefficient or lags (peak: p=0.16; lags: p=0.23), or genotype MK801 interaction effect (peak: p=0.79; lags: p =0.46), suggesting that stimulus-response decorrelation in rTg4510 neurons was not dependent on NMDAR function *(Figure 8C-D&F)*. Thus, whilst synaptic responses in WT neurons were faithfully temporally modulated by 5 Hz whisker deflection, encoding of this rhythmic sensory input was substantially degraded in TG neurons.

**Figure 8.**
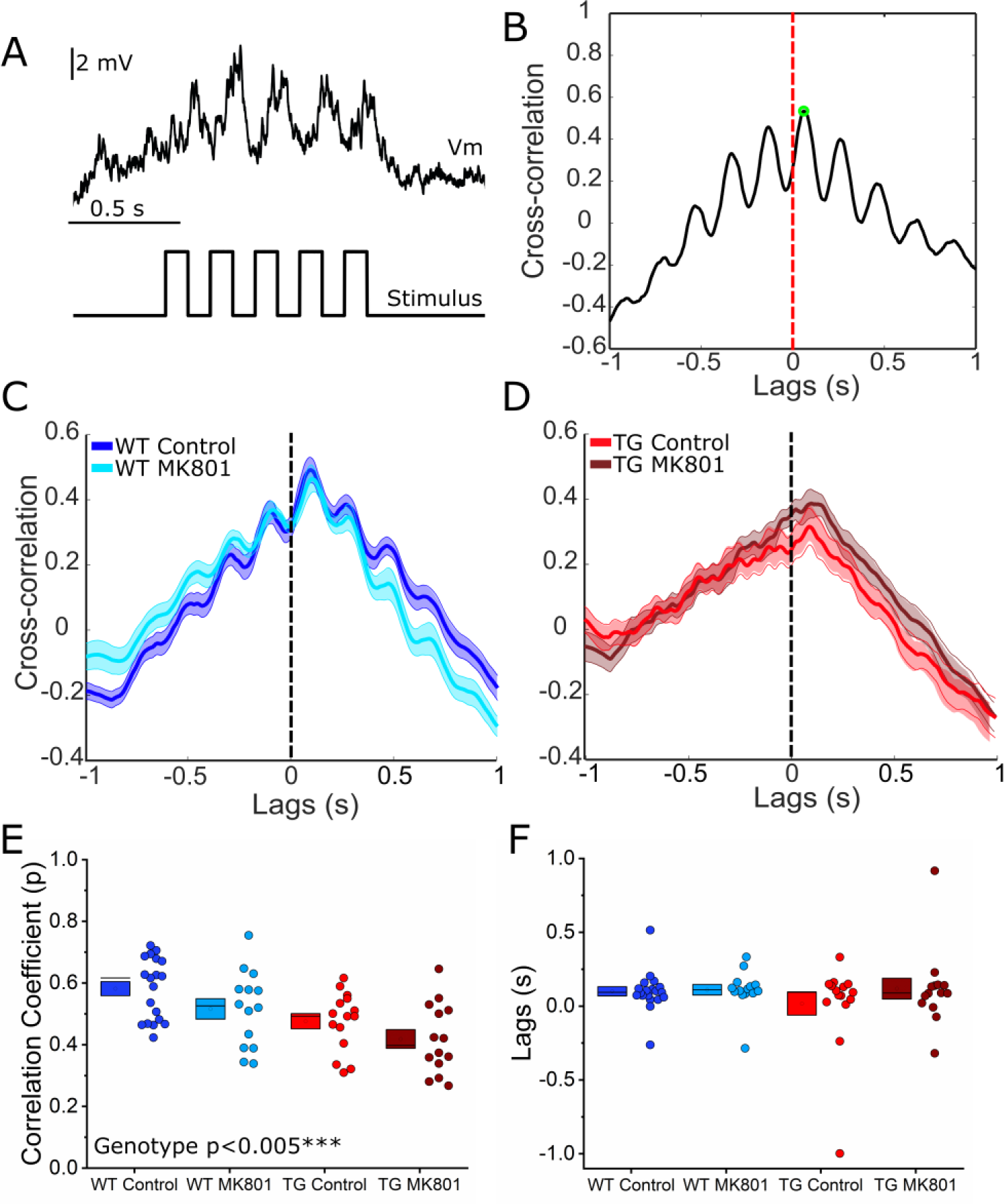
The cross-correlation between 5 Hz sensory input and the evoked synaptic response is reduced in rTg4510 cortical pyramidal neurons at 5.5 M. **A)** An example voltage trace (top) illustrating the synaptic response to 5 Hz whisker stimulation (bottom) in a WT putative somatosensory cortex pyramidal neuron. **B)** Example cross-correlation between membrane voltage and whisker stimulation traces for the neuron illustrated in (A). The peak of the cross-correlation is annotated by a green circle; 0 lags is annotated by a red dashed line. **C)** Average cross-correlation for WT neurons, with or without MK801 treatment. Thick lines represent the mean, with shading representing SEM. **D)** Average cross-correlation for TG neurons, with or without MK801 treatment. Thick lines represent the mean, with shading representing SEM. **E)** Box and dot plots showing the peak cross-correlation coefficient for each neuron. **F)** Box and dot plots showing the lag of the peak correlation coefficient for each neuron. Statistics: GLMM, main significant effects labelled on the graph, with statistical significance denoted by asterisks (p<0.005***).

## Discussion

We have investigated perturbations to cortical synapses in the early phases of tauopathy. While at the tissue level, loss of synaptic proteins was linked to disease progression, we found subtleties in how these effects link to synaptic function and changes in neuronal morphology. Specifically, our data suggests that early tauopathy is associated with a shift in the spatial and temporal organisation of synaptic inputs to cortical pyramidal neurons. As a result, the ability of these neurons to faithfully encode temporally coordinated synaptic signals is degraded, suggesting a mechanism that may underlie early symptoms of tauopathy.

One point to consider is how these results from a single rodent model that recapitulates some aspects of dementia-like pathology, generalises to human tauopathies. In particular, the rTg4510 mouse model is now known to exhibit off-target mutations around the site of MAPT^P301L^ transgene insertion that may contribute to pathological phenotypes.^78,79^ However, studies that have used doxycycline to suppress transgene expression at different developmental age points in rTg4510 mice have showed a range of tauopathy-driven cellular and synaptic phenotypes.^40,80^ Overall, the rTg4510 mouse is one of only a few well-characterised models that displays robust tau pathogenesis and neurodegeneration. Nonetheless, studies examining neuronal morphology and synaptic alterations in other models of early and progressed tauopathy, such as those with mutations expressed under endogenous promoters,^81^ will be important comparators to this work.

Biochemical assessment of cortical synaptosomes showed a profound decrease in PSD95 – a key component of excitatory postsynapses – in rTg4510 mice (*Figure 1*). The magnitude of this effect was not mirrored by synaptophysin – a key presynaptic protein – which was subtly reduced in rTg4510 mice (*Figure 1*). This suggests a particular vulnerability of postsynapses compared to presynaptic compartments to tauopathy. Previous longitudinal imaging of axonal boutons and dendritic spines (synonymous with pre-and post-synaptic components, respectively) within similarly aged (5.5 M) rTg4510 mice showed increased bouton stability accompanied by increased spine turnover.^42,43^ Divergence of functional pathology in pre- and post-synaptic compartments may be a feature of early tauopathy. Within the postsynapse, PSD95 anchors glutamatergic receptors, promoting synaptic stability.^82,83^ Therefore, decreased stability of dendritic spines could be due to lower levels of PSD95. Many studies have reported decreased levels of pre- and post-synaptic proteins and structures in late, degenerative stages in rTg4510 mice, and in other models of tauopathy.^23,35,84^ This is usually thought to reflect a generalised synapse loss that accompanies frank neurodegeneration. The reduced expression of PSD95 and synaptophysin we report in 5.5 M rTg4510 mice may reflect an early loss of particularly vulnerable synapses, or a period of neuronal and synaptic pathophysiology. Our biochemical, structural, and electrophysiological data suggest both occur.

We observed a substantial decrease in several AMPAR and NMDAR subunits (specifically, GluA2, GluA3, and GluN1) in somatosensory cortex samples from rTg4510 mice (*Figure 2*). We were therefore surprised to find no change in the NMDA:AMPA receptor EPSC ratio in electrophysiological recordings of L4 input to L2/3 pyramidal neurons (*Figure 4*). We also observed no change in PPR as a measure of short-term synaptic plasticity in this pathway, suggesting that presynaptic release is likely unaffected at these synapses in early tauopathy (*Figure 3*). It is possible that intracolumnar L4-to-L2/3 synapses were preserved, whilst effects on other synapses in the same cortical region caused changes in synaptic protein expression in rTg4510 mice, implying that specific subpopulations of synapses could be more vulnerable to tauopathy than others. Indeed, we observed changes to the morphology of dendrites proximal to the soma of 5.5 M rTg4510 pyramidal neurons, suggesting that tauopathy can drive alterations in specific parts of the neuron (*Figure 5*). In degenerative stages of pathology (∼7-13 months old) in rTg4510 mice and other murine tauopathy models, there is evidence of dendritic atrophy in cortical pyramidal neurons reminiscent of that found in Alzheimer’s disease post-mortem brain tissue.^26,30,31,34,35^ Notably, however, it was shown that some neurons appear spared from degeneration.^26^ The dendritic changes we observed in 5.5M rTg4510 neurons were not associated with overt signs of cellular atrophy, suggesting that dendritic remodelling could represent an early homeostatic compensation for altered synaptic input. Increased branching proximal to the soma could lead to an altered balance of inputs from different anatomically distributed synaptic pathways that target specific parts of the dendritic tree.^67,68,85,86^ For example, local and sub-cortical inputs tend to cluster on basal or proximal dendrites, whereas apical dendrites tend to receive long-range inputs.^67,68,70,85,86^ If additional synapses are formed onto the more complex proximal branches of rTg4510 neurons, we would anticipate a shift in the spatial integration of synaptic input that might alter neuronal response properties.

To investigate how changes in synaptic receptor expression and dendritic structure contribute to changes in signal processing, *in vivo* whole-cell patch clamp recordings were made during whisker stimulation to evoke physiological multimodal synaptic responses resulting from integration of local (intracolumnar) and long-range (intercolumnar) synaptic input. These experiments revealed a functional impact of early tauopathy on synaptic encoding. Depolarisation evoked by the first whisker deflection was increased in rTg4510 neurons, revealing altered synaptic input (*Figure 6*). This brief depolarisation is likely dominated by direct thalamocortical input, which preferentially targets the proximal basal dendrites that we found were more complex in L2/3 rTg4510 neurons (*Figure 5*).^70^ It is possible that thalamocortical axons form more synapses onto these proximal dendrites, causing increased depolarisation. The initial EPSP was, in part, mediated by NMDARs because its peak was decreased by MK801 (*Figure 6*). However, we did not find any difference in NMDAR contribution between WT and rTg4510 neurons to these EPSPs.

Rhythmic (5 Hz) trains of whisker deflections generated sustained postsynaptic depolarisation in WT and rTg4510 neurons (*Figure 7*). However, the shape of the response to rhythmic multi-whisker stimulation was markedly different. Whereas WT neurons exhibited frequency modulated responses to rhythmic input, this was blunted in rTg4510 neurons (*Figure 8*). The PSP waveform elicited by this protocol likely reflects the integration of excitatory and inhibitory synaptic potentials arising from multiple intracolumnar and intercolumnar pathways.^54,73,75^ We speculate that changes in GABAergic inhibition onto rTg4510 neurons could potentially explain this change in the response dynamics. Notably, local inhibition evokes inhibitory postsynaptic potentials 10-20 ms post whisker stimulation^55^, and stimulation of multiple whiskers activates excitatory and inhibitory inputs in multiple cortical columns that contribute to the compound PSP.^73^ Deficits in synaptic protein expression – like those we have observed (*Figure 1 & Figure* 2) – in GABAergic interneurons, may reduce their ability to be recruited by sensory input. Evidence to support this idea comes from observation of reduced synapse density in interneurons during early tauopathy in the rTg4510 model.^87^ Consistent with this, the timing of interneuron activity is disrupted during sharp wave-ripple oscillations in the hippocampus of rTg4510 mice,^36^ implying that disrupted timing of GABAergic input may hinder entrainment of rTg4510 neurons to rhythmic input. Therefore, alterations in coordinated synaptically-driven activity are not only observed in the somatosensory cortex and may be a generalised mechanism of early neural circuit disruption in tauopathy.

## Conclusions

Our research has identified significant reductions in glutamatergic receptor and synaptic marker expression during early tauopathy. These changes occurred alongside altered dendritic structure in cortical pyramidal neurons, which was potentially linked to changes in synaptic function, particularly entrainment of synaptic responses to rhythmic multimodal input. Our findings suggest that synaptic alterations in early tauopathy likely manifest through disrupted coordination of neural network activity. Understanding how such changes precipitate into altered signal processing may be critical to preventing symptoms in tauopathy-associated dementia.

## Abbreviations

aCSF: artificial cerebrospinal fluid
AD: Alzheimer’s disease
AMPAR: AMPA receptor
AP: action potential
AUC: area under the curve
EPSC: excitatory postsynaptic current
GLMM: Generalised linear mixed model
ICC: intraclass coefficient
L2/3: layer 2/3
L4: layer 4
M: Month
NMDAR: NMDA receptor
PB: phosphate buffer
PBS: phosphate buffered saline
PBS-T: phosphate buffered saline with 0.05% tween
PBS-T: A 0.1M PBS containing 0.3% Triton-X and 0.05% sodium azide
PBS-T-M: phosphate buffered saline with 0.05% tween and 5% dried milk
PPR: paired pulse ratio
PSP: postsynaptic potential
RT: Room temperature
rTg4510: regulatable Tg4510
SEM: standard error of the mean
TG: transgenic
TPP: 50 mM Tris containing phosphatase and protease inhibitors.
Vm: membrane voltage
WT: wild-type

## Acknowledgements

We thank Jean-Sebastien Jouhanneau, Leiron Ferrarese and James Poulet for training SM to perform *in vivo* whole-cell patch clamp electrophysiology. We also thank Charlotte Dunbar and Claire Hardy for training SM to perform western blots.

