## Supplementary materials for "Synaptic alterations associated with disrupted sensory encoding in a mouse model of tauopathy"

#### *Table S1. List of primary and secondary antibodies used in synaptosome experiments*

| **Antibody** | **Target** | **Species** | **Dilution used** | **Supplier** | **Catalogue No.** | **Lot No.** |
|---|---|---|---|---|---|---|
| GAPDH | Housekeeping proteins | Mouse | 1:6000 | Invitrogen | AM4300 | Multiple |
| AT-8 | Tau | Mouse | 1:1000 | Peter Davies | - | - |
| CP27 | Tau | Mouse | 1:1000 | Peter Davies | - | - |
| PSD95 | PSD95 | Mouse | 1:1000 | BD | 610495 | 8128742 |
| Synaptophysin | Synaptic vesicles | Mouse | 1:1000 | Abcam | Ab8049 (SY38) | GR3280108-1 |
| GluR1 (AMPA) [EPR5479] | AMPA receptor, subunit 1 | Rabbit | 1:1000 | Abcam | Ab109450 | GR3241245-1 |
| Anti-GluA2/GluR2 Glutamate Receptor Clone L21/32 | AMPA receptor, subunit 2 | Mouse | 1:1000 | NeuroMab | 75-002 | 472-IJU-17 |
| GluR3 | AMPA receptor, subunit 3 | Mouse | 1:1000 | Invitrogen | 32-0400 | RH240594 |
| NMDAR1 Monoclonal Antibody (54.1) | NMDA receptor, subunit 1 | Mouse | 1:1000 | Invitrogen | 32-0500 | UH286597 |
| Anti-NR2A, M264-10ug | NMDA receptor, subunit 2A | Rabbit | 1:1000 | Sigma | 1002457527 | MKCC8197 |
| NMDAR2B | NMDA receptor, subunit 2B | Mouse | 1:1000 | BD | 610416 | 8159860 |
| GluN2C clone N422/18 | NMDA receptor, subunit 2C | Mouse | 1:1000 | NeuroMab | 75-411 | 455-6JD-37 |

| | | | | | | |
|---|---|---|---|---|---|---|
| NR2D | NMDA receptor, subunit 2D | Rabbit | 1:1000 | Abcam | Ab35448 | 851239 |
| HRP linked anti-mouse IgG | Mouse raised immunogens | - | 1:20000 | Cell Signalling | 7076S | Multiple |
| ECL anti-rabbit IgG HRP linked | Rabbit raised immunogens | - | 1:10000 | GE | NA934V | Multiple |

#### Western Blot Images

The ladder is labelled with corresponding molecular weights.

Key for first line labels: SCTX = Somatosensory cortex; Blank = Blank lane; P301S = Positive Control; L = Ladder.

Key for second line labels: Number is sample number; B = Blank; Ctrl = Positive Control; L = Ladder; Blue = 5.5M WT; Green = 7.5M WT; Orange = 5.5M TG; Grey = 7.5M TG.

Third line labels are the lane number.

Exclusions due to incomplete or undetectable bands in either the test antibody or GAPDH.

Synaptophysin: Samples 35 & 36.

PSD95: Samples 35 & 36.

GluA1: Sample 36.

GluA2: Sample 1, 35, 36.

GluA3: Sample 34.

GluN1: None.

### Synaptophysin

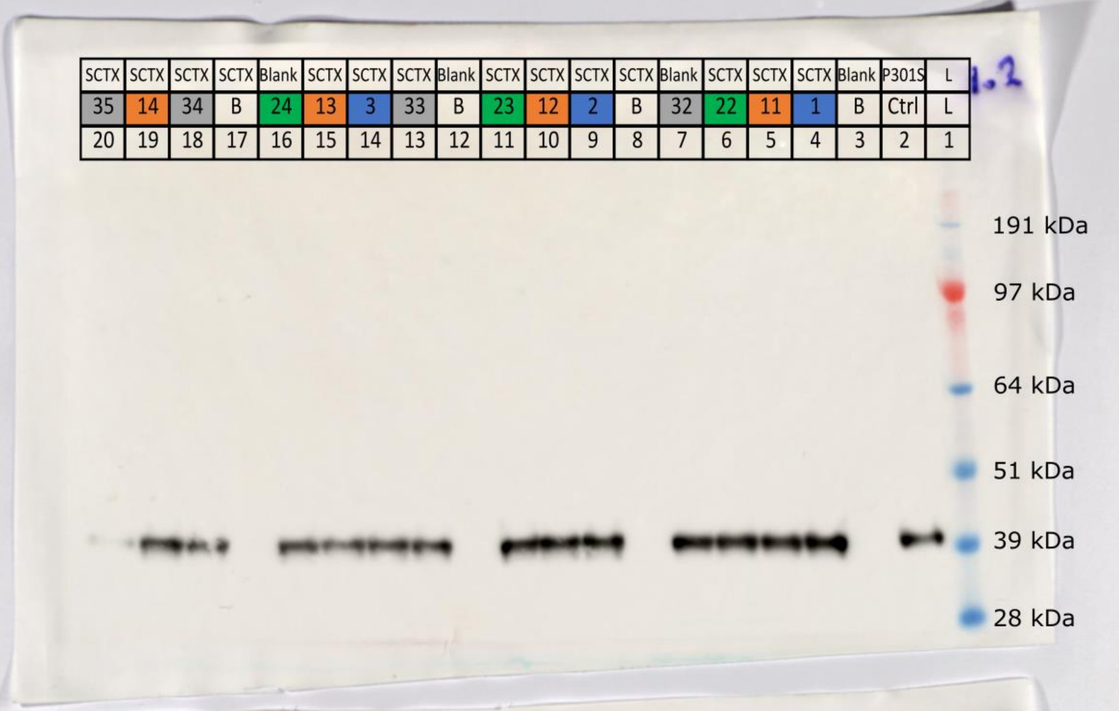


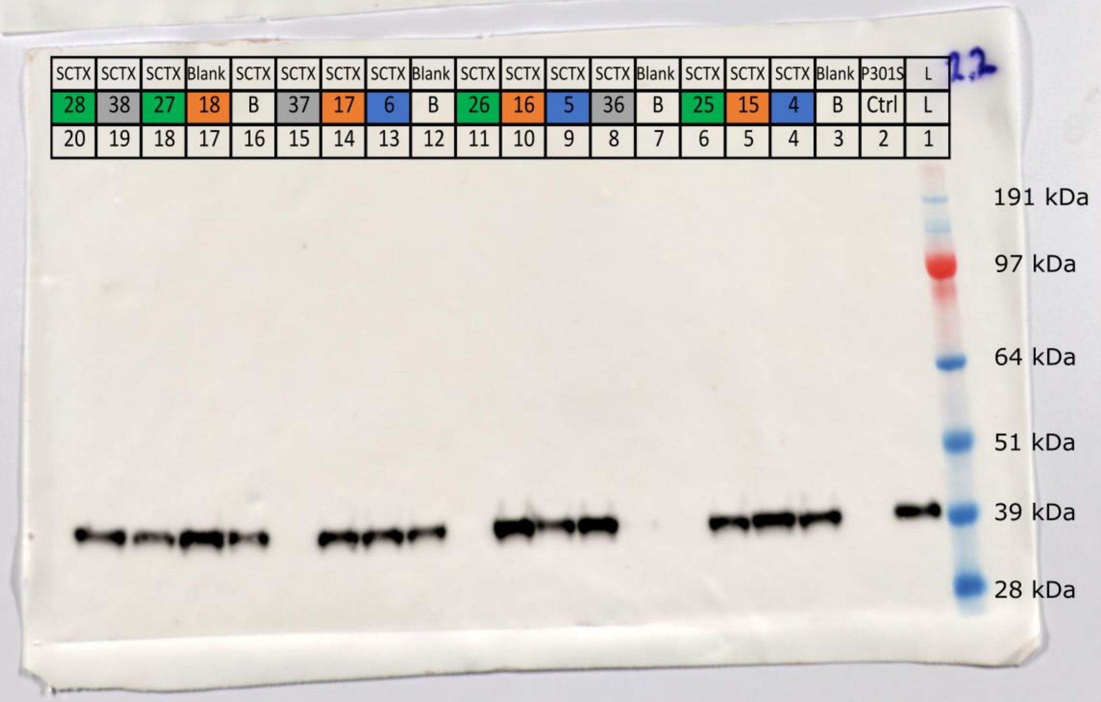

Synaptophysin

| | | | | | | | | | | | | | | | | | | | |
|---|---|---|---|---|---|---|---|---|---|---|---|---|---|---|---|---|---|---|---|
| SCTX | SCTX | SCTX | SCTX | Blank | SCTX | SCTX | SCTX | Blank | SCTX | SCTX | SCTX | Blank | SCTX | SCTX | SCTX | SCTX | Blank | P301S | L |
| 31 | 42 | 21 | 10 | B | 41 | 9 | 30 | B | 8 | 40 | 20 | B | 39 | 29 | 19 | 7 | B | Ctrl | L |
| 20 | 19 | 18 | 17 | 16 | 15 | 14 | 13 | 12 | 11 | 10 | 9 | 8 | 7 | 6 | 5 | 4 | 3 | 2 | 1 |

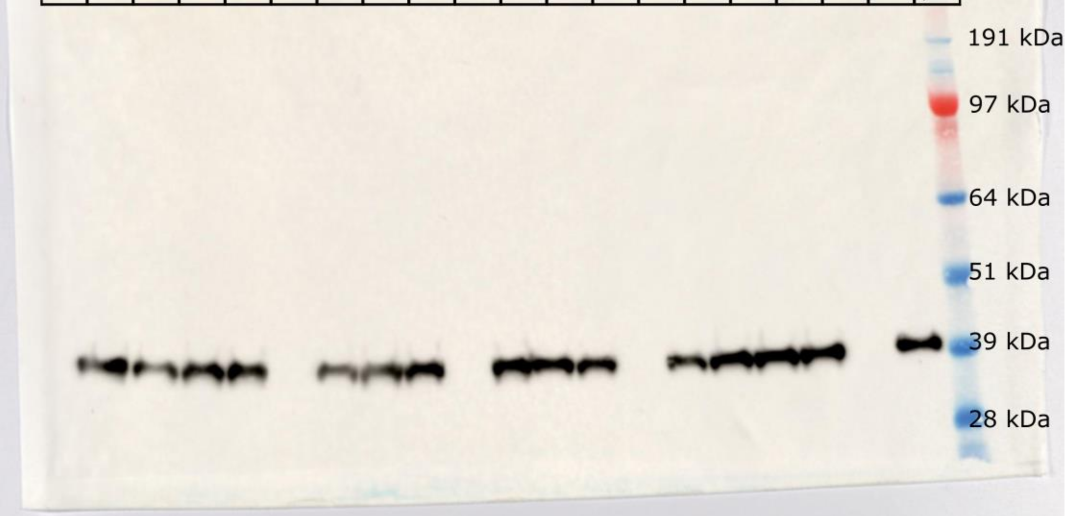

### Synaptophysin - GAPDH Control

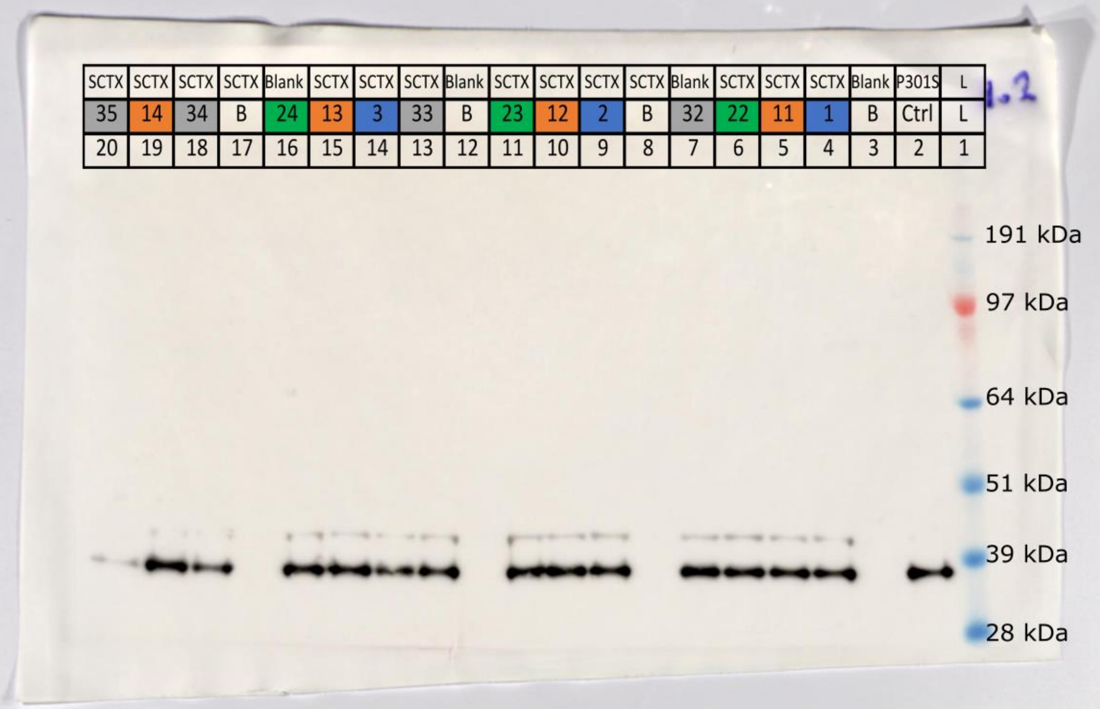


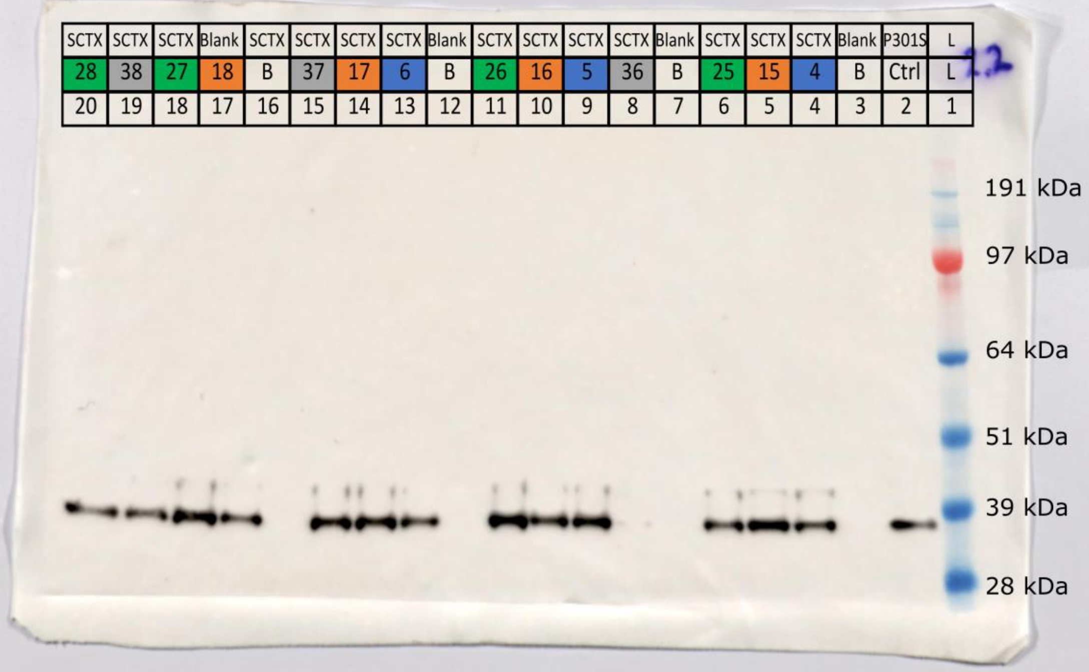

### Synaptophysin - GAPDH Control

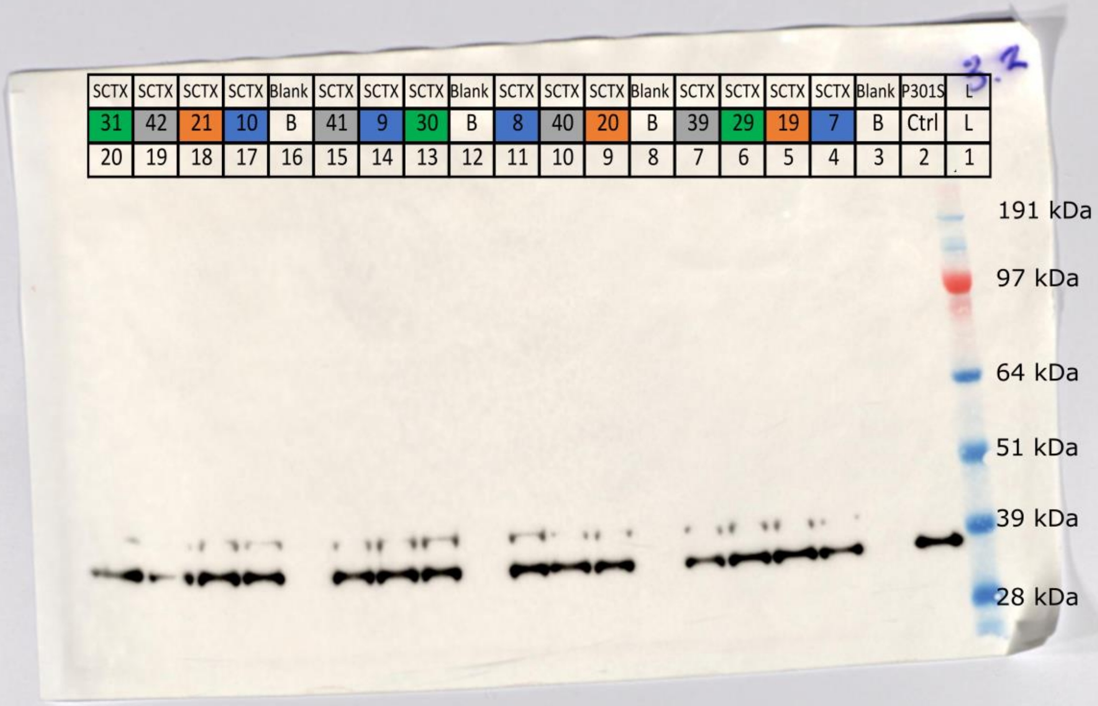

### PSD95

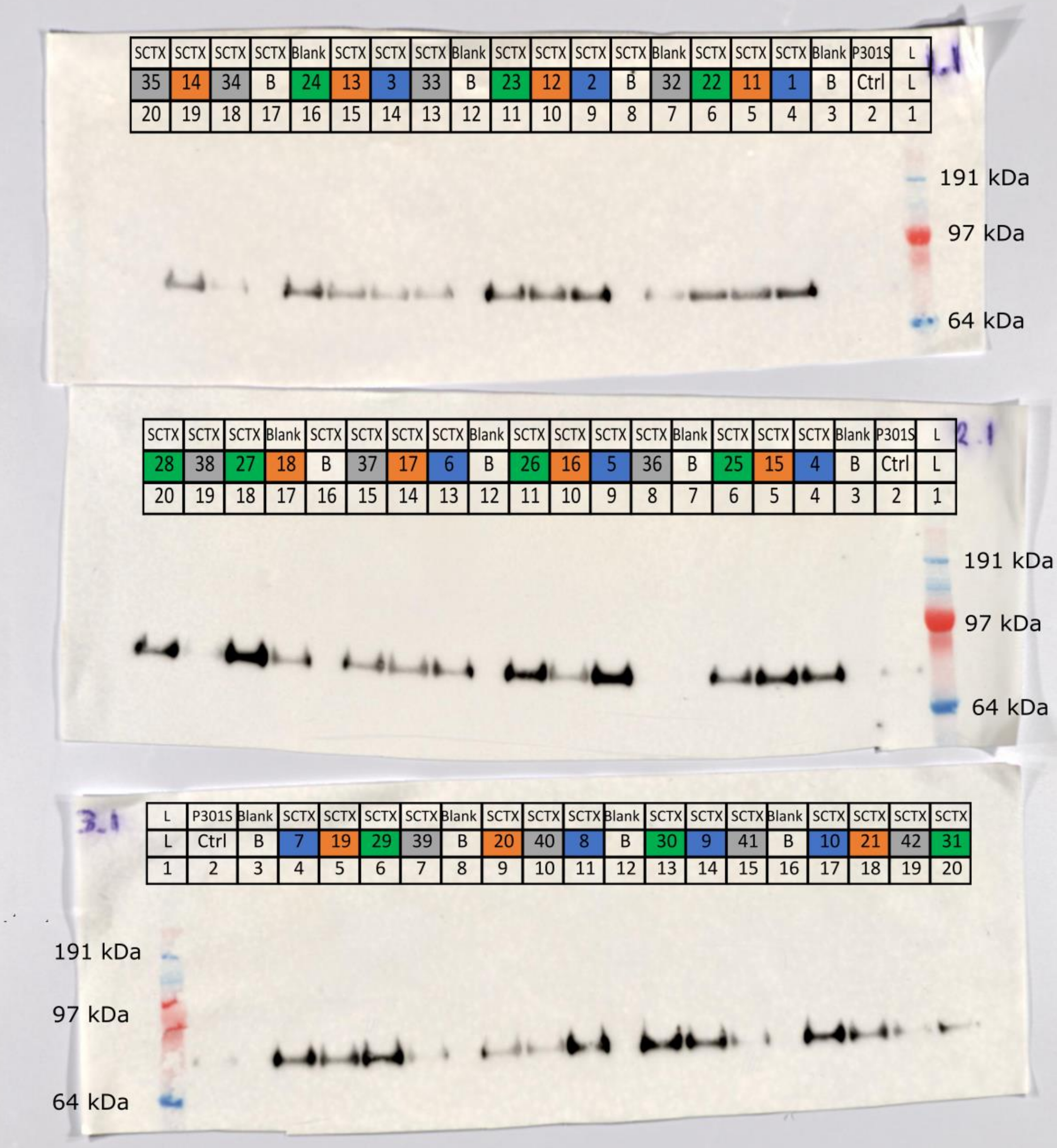
1.1
SCTX SCTX SCTX SCTX Blank SCTX SCTX SCTX Blank SCTX SCTX SCTX SCTX Blank SCTX SCTX SCTX Blank P301S L
35 14 34 B 24 13 3 33 B 23 12 2 B 32 22 11 1 B Ctrl L
20 19 18 17 16 15 14 13 12 11 10 9 8 7 6 5 4 3 2 1
191 kDa
97 kDa
64 kDa
2.1
SCTX SCTX SCTX Blank SCTX SCTX SCTX SCTX Blank SCTX SCTX SCTX SCTX Blank SCTX SCTX SCTX Blank P301S L
28 38 27 18 B 37 17 6 B 26 16 5 36 B 25 15 4 B Ctrl L
20 19 18 17 16 15 14 13 12 11 10 9 8 7 6 5 4 3 2 1
191 kDa
97 kDa
64 kDa
3.1
L P301S Blank SCTX SCTX SCTX SCTX Blank SCTX SCTX SCTX Blank SCTX SCTX SCTX Blank SCTX SCTX SCTX SCTX
L Ctrl B 7 19 29 39 B 20 40 8 B 30 9 41 B 10 21 42 31
1 2 3 4 5 6 7 8 9 10 11 12 13 14 15 16 17 18 19 20
191 kDa
97 kDa
64 kDa

### PSD95 - GAPDH Control

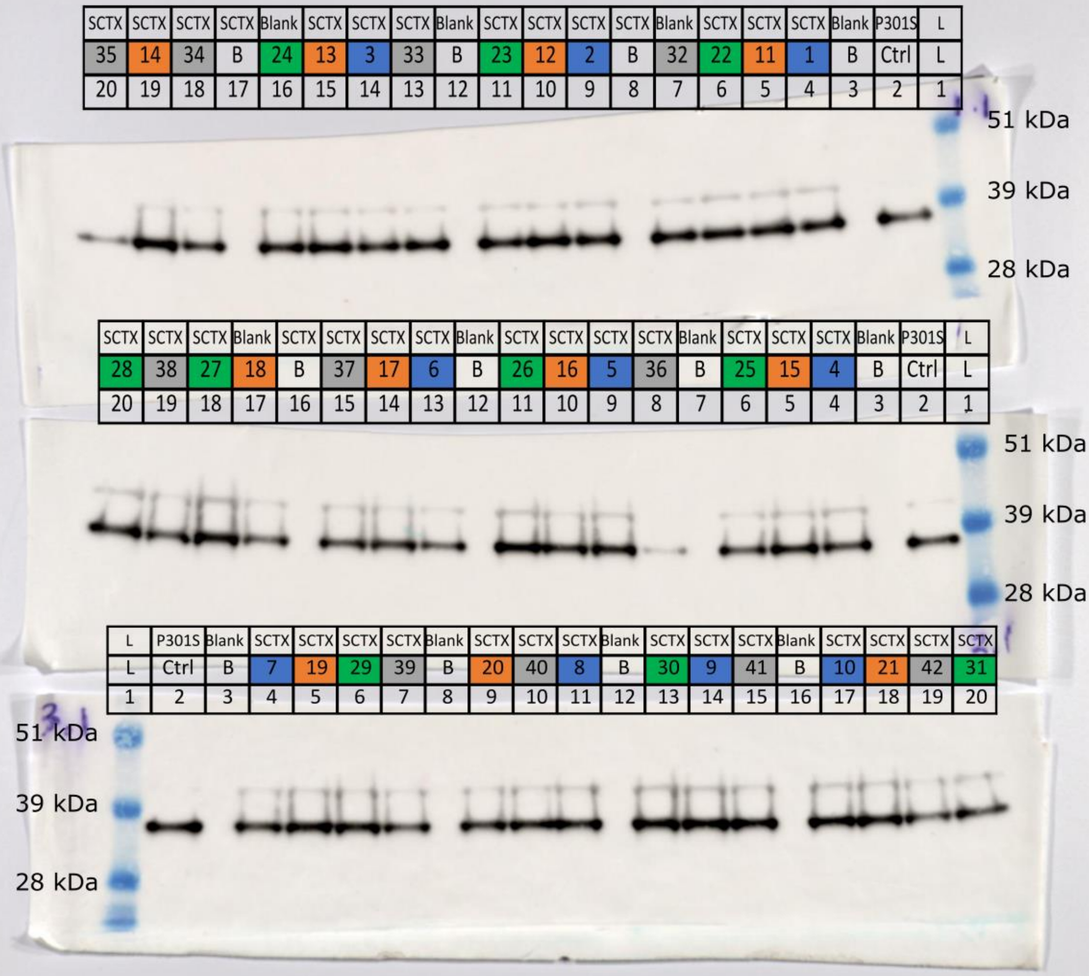

### GluA1

1.3

| | | | | | | | | | | | | | | | | | | | |
|---|---|---|---|---|---|---|---|---|---|---|---|---|---|---|---|---|---|---|---|
| L | P301S | Blank | SCTX | SCTX | SCTX | Blank | SCTX | SCTX | SCTX | SCTX | Blank | SCTX | SCTX | SCTX | Blank | SCTX | SCTX | SCTX | SCTX |
| L | Ctrl | B | 1 | 11 | 22 | 32 | B | 2 | 12 | 23 | B | 33 | 3 | 13 | 24 | B | 34 | 14 | 35 |
| 1 | 2 | 3 | 4 | 5 | 6 | 7 | 8 | 9 | 10 | 11 | 12 | 13 | 14 | 15 | 16 | 17 | 18 | 19 | 20 |

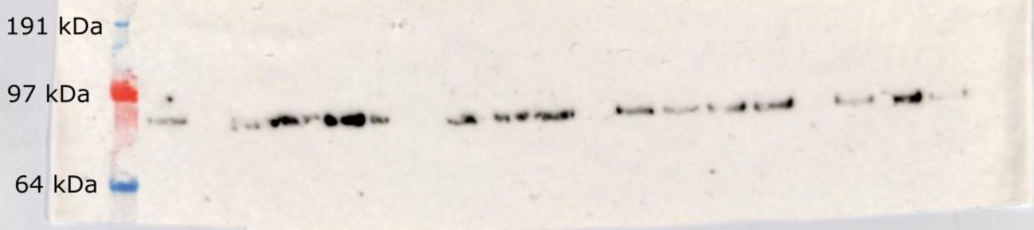


2.3

| | | | | | | | | | | | | | | | | | | | |
|---|---|---|---|---|---|---|---|---|---|---|---|---|---|---|---|---|---|---|---|
| L | P301S | Blank | SCTX | SCTX | SCTX | Blank | SCTX | SCTX | SCTX | SCTX | Blank | SCTX | SCTX | SCTX | SCTX | Blank | SCTX | SCTX | SCTX |
| L | Ctrl | B | 4 | 15 | 25 | B | 36 | 5 | 16 | 26 | B | 6 | 17 | 37 | B | 18 | 27 | 38 | 28 |
| 1 | 2 | 3 | 4 | 5 | 6 | 7 | 8 | 9 | 10 | 11 | 12 | 13 | 14 | 15 | 16 | 17 | 18 | 19 | 20 |

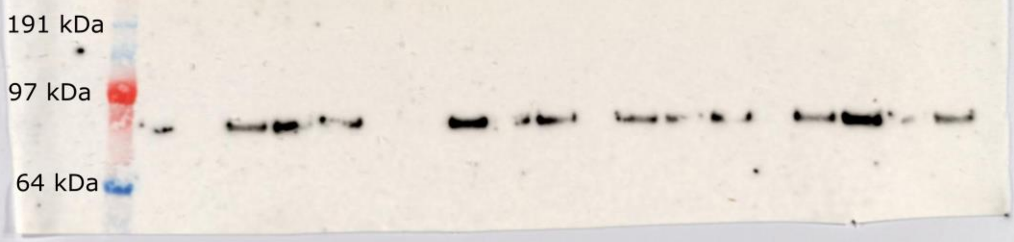


3.3

| | | | | | | | | | | | | | | | | | | | |
|---|---|---|---|---|---|---|---|---|---|---|---|---|---|---|---|---|---|---|---|
| L | P301S | Blank | SCTX | SCTX | SCTX | SCTX | Blank | SCTX | SCTX | SCTX | Blank | SCTX | SCTX | SCTX | Blank | SCTX | SCTX | SCTX | SCTX |
| L | Ctrl | B | 7 | 19 | 29 | 39 | B | 20 | 40 | 8 | B | 30 | 9 | 41 | B | 10 | 21 | 42 | 31 |
| 1 | 2 | 3 | 4 | 5 | 6 | 7 | 8 | 9 | 10 | 11 | 12 | 13 | 14 | 15 | 16 | 17 | 18 | 19 | 20 |

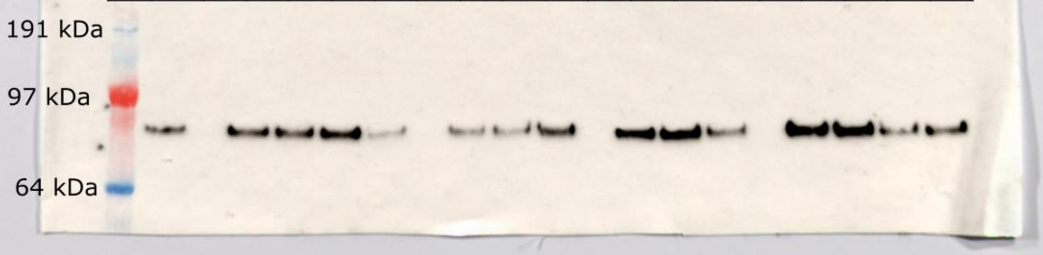

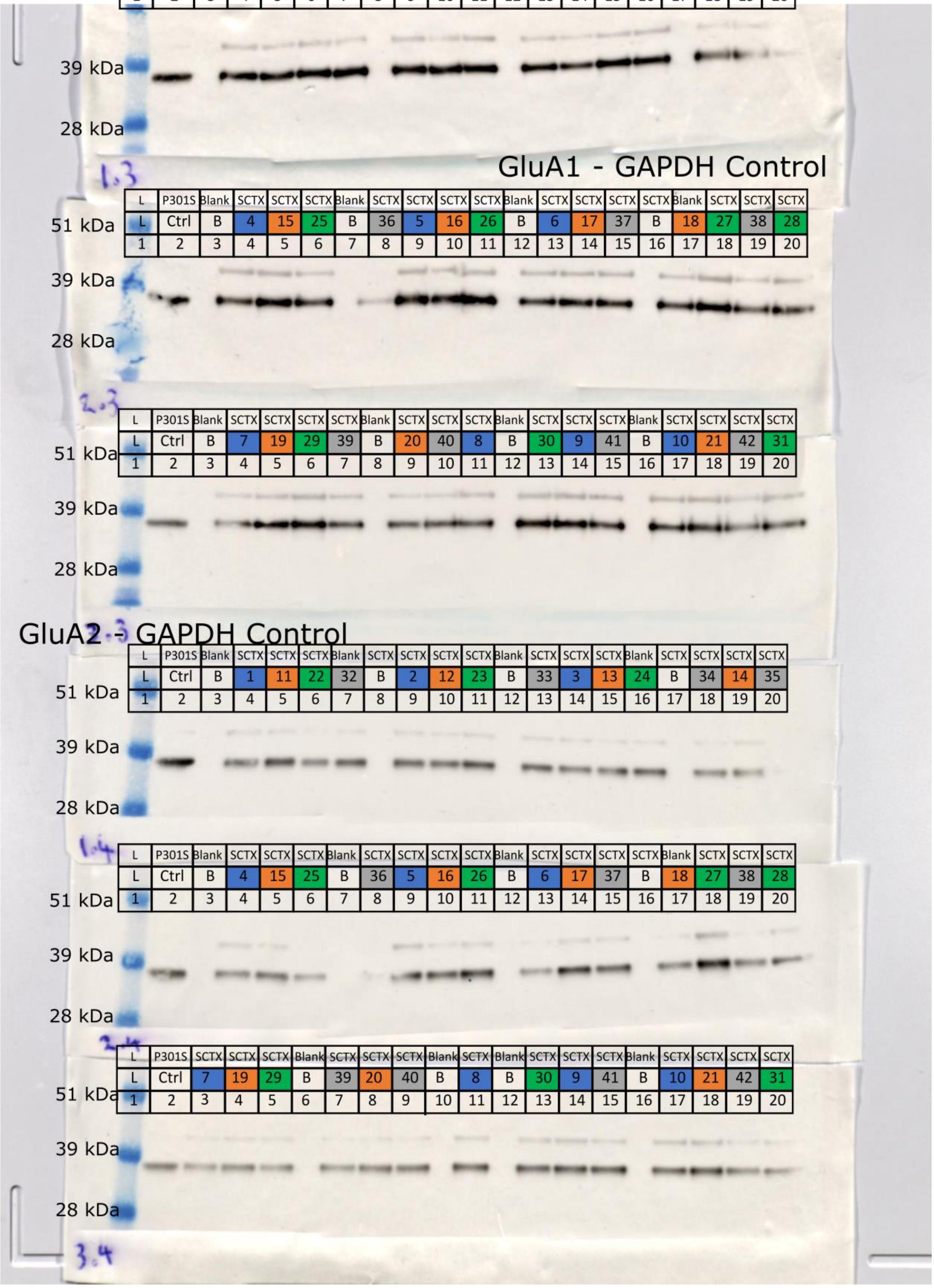


| L | P301S | Blank | SCTX | SCTX | SCTX | Blank | SCTX | SCTX | SCTX | SCTX | Blank | SCTX | SCTX | SCTX | Blank | SCTX | SCTX | SCTX | SCTX |
|---|---|---|---|---|---|---|---|---|---|---|---|---|---|---|---|---|---|---|---|
| L | Ctrl | B | 1 | 11 | 22 | 32 | B | 2 | 12 | 23 | B | 33 | 3 | 13 | 24 | B | 34 | 14 | 35 |
| 1 | 2 | 3 | 4 | 5 | 6 | 7 | 8 | 9 | 10 | 11 | 12 | 13 | 14 | 15 | 16 | 17 | 18 | 19 | 20 |

#### GluA1 - GAPDH Control

| L | P301S | Blank | SCTX | SCTX | SCTX | Blank | SCTX | SCTX | SCTX | SCTX | Blank | SCTX | SCTX | SCTX | SCTX | Blank | SCTX | SCTX | SCTX |
|---|---|---|---|---|---|---|---|---|---|---|---|---|---|---|---|---|---|---|---|
| L | Ctrl | B | 4 | 15 | 25 | B | 36 | 5 | 16 | 26 | B | 6 | 17 | 37 | B | 18 | 27 | 38 | 28 |
| 1 | 2 | 3 | 4 | 5 | 6 | 7 | 8 | 9 | 10 | 11 | 12 | 13 | 14 | 15 | 16 | 17 | 18 | 19 | 20 |

| L | P301S | Blank | SCTX | SCTX | SCTX | SCTX | Blank | SCTX | SCTX | SCTX | Blank | SCTX | SCTX | SCTX | Blank | SCTX | SCTX | SCTX | SCTX |
|---|---|---|---|---|---|---|---|---|---|---|---|---|---|---|---|---|---|---|---|
| L | Ctrl | B | 7 | 19 | 29 | 39 | B | 20 | 40 | 8 | B | 30 | 9 | 41 | B | 10 | 21 | 42 | 31 |
| 1 | 2 | 3 | 4 | 5 | 6 | 7 | 8 | 9 | 10 | 11 | 12 | 13 | 14 | 15 | 16 | 17 | 18 | 19 | 20 |

#### GluA2 - GAPDH Control

| L | P301S | Blank | SCTX | SCTX | SCTX | Blank | SCTX | SCTX | SCTX | SCTX | Blank | SCTX | SCTX | SCTX | Blank | SCTX | SCTX | SCTX | SCTX |
|---|---|---|---|---|---|---|---|---|---|---|---|---|---|---|---|---|---|---|---|
| L | Ctrl | B | 1 | 11 | 22 | 32 | B | 2 | 12 | 23 | B | 33 | 3 | 13 | 24 | B | 34 | 14 | 35 |
| 1 | 2 | 3 | 4 | 5 | 6 | 7 | 8 | 9 | 10 | 11 | 12 | 13 | 14 | 15 | 16 | 17 | 18 | 19 | 20 |

| L | P301S | Blank | SCTX | SCTX | SCTX | Blank | SCTX | SCTX | SCTX | SCTX | Blank | SCTX | SCTX | SCTX | SCTX | Blank | SCTX | SCTX | SCTX |
|---|---|---|---|---|---|---|---|---|---|---|---|---|---|---|---|---|---|---|---|
| L | Ctrl | B | 4 | 15 | 25 | B | 36 | 5 | 16 | 26 | B | 6 | 17 | 37 | B | 18 | 27 | 38 | 28 |
| 1 | 2 | 3 | 4 | 5 | 6 | 7 | 8 | 9 | 10 | 11 | 12 | 13 | 14 | 15 | 16 | 17 | 18 | 19 | 20 |

| L | P301S | SCTX | SCTX | SCTX | Blank | SCTX | SCTX | SCTX | Blank | SCTX | Blank | SCTX | SCTX | SCTX | Blank | SCTX | SCTX | SCTX | SCTX |
|---|---|---|---|---|---|---|---|---|---|---|---|---|---|---|---|---|---|---|---|
| L | Ctrl | 7 | 19 | 29 | B | 39 | 20 | 40 | B | 8 | B | 30 | 9 | 41 | B | 10 | 21 | 42 | 31 |
| 1 | 2 | 3 | 4 | 5 | 6 | 7 | 8 | 9 | 10 | 11 | 12 | 13 | 14 | 15 | 16 | 17 | 18 | 19 | 20 |

### GluA2

1.4

| | | | | | | | | | | | | | | | | | | | |
|---|---|---|---|---|---|---|---|---|---|---|---|---|---|---|---|---|---|---|---|
| L | P301S | Blank | SCTX | SCTX | SCTX | Blank | SCTX | SCTX | SCTX | SCTX | Blank | SCTX | SCTX | SCTX | Blank | SCTX | SCTX | SCTX | SCTX |
| L | Ctrl | B | 1 | 11 | 22 | 32 | B | 2 | 12 | 23 | B | 33 | 3 | 13 | 24 | B | 34 | 14 | 35 |
| 1 | 2 | 3 | 4 | 5 | 6 | 7 | 8 | 9 | 10 | 11 | 12 | 13 | 14 | 15 | 16 | 17 | 18 | 19 | 20 |

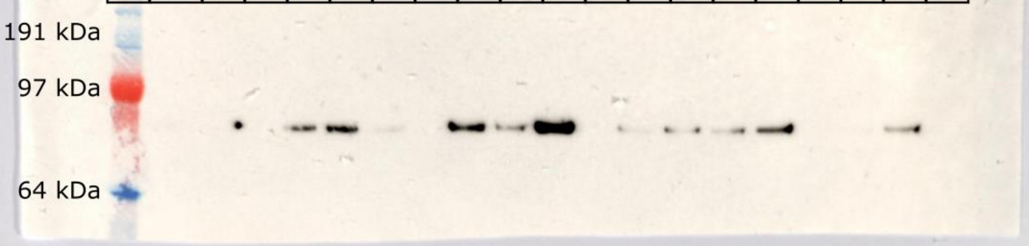


2.4

| | | | | | | | | | | | | | | | | | | | |
|---|---|---|---|---|---|---|---|---|---|---|---|---|---|---|---|---|---|---|---|
| L | P301S | Blank | SCTX | SCTX | SCTX | Blank | SCTX | SCTX | SCTX | SCTX | Blank | SCTX | SCTX | SCTX | SCTX | Blank | SCTX | SCTX | SCTX |
| L | Ctrl | B | 4 | 15 | 25 | B | 36 | 5 | 16 | 26 | B | 6 | 17 | 37 | B | 18 | 27 | 38 | 28 |
| 1 | 2 | 3 | 4 | 5 | 6 | 7 | 8 | 9 | 10 | 11 | 12 | 13 | 14 | 15 | 16 | 17 | 18 | 19 | 20 |

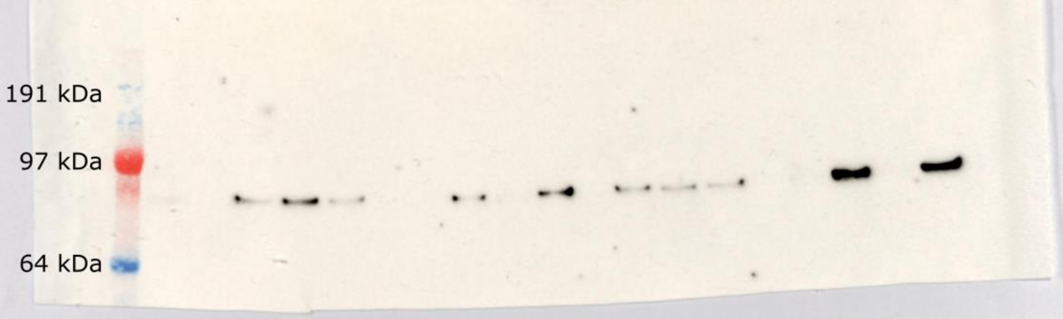


3.4

| | | | | | | | | | | | | | | | | | | | |
|---|---|---|---|---|---|---|---|---|---|---|---|---|---|---|---|---|---|---|---|
| L | P301S | SCTX | SCTX | SCTX | Blank | SCTX | SCTX | SCTX | Blank | SCTX | Blank | SCTX | SCTX | SCTX | Blank | SCTX | SCTX | SCTX | SCTX |
| L | Ctrl | 7 | 19 | 29 | B | 39 | 20 | 40 | B | 8 | B | 30 | 9 | 41 | B | 10 | 21 | 42 | 31 |
| 1 | 2 | 3 | 4 | 5 | 6 | 7 | 8 | 9 | 10 | 11 | 12 | 13 | 14 | 15 | 16 | 17 | 18 | 19 | 20 |

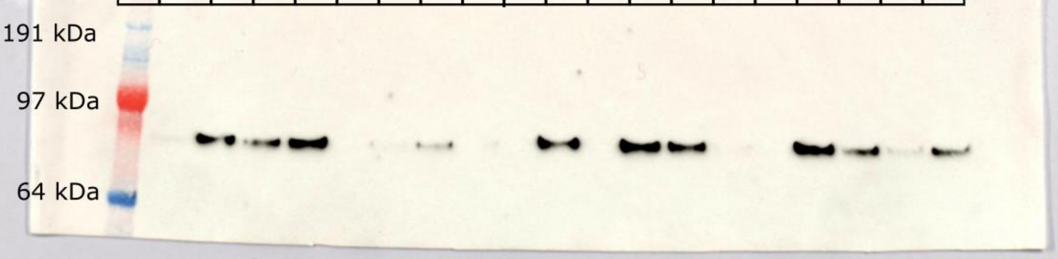

### GluA3

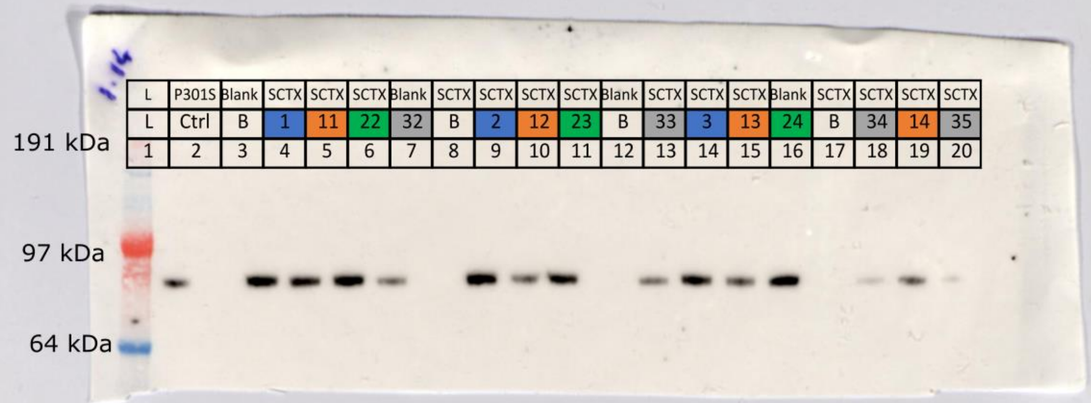

1.14

| L | P301S | Blank | SCTX | SCTX | SCTX | Blank | SCTX | SCTX | SCTX | SCTX | Blank | SCTX | SCTX | SCTX | Blank | SCTX | SCTX | SCTX | SCTX |
|---|---|---|---|---|---|---|---|---|---|---|---|---|---|---|---|---|---|---|---|
| L | Ctrl | B | 1 | 11 | 22 | 32 | B | 2 | 12 | 23 | B | 33 | 3 | 13 | 24 | B | 34 | 14 | 35 |
| 1 | 2 | 3 | 4 | 5 | 6 | 7 | 8 | 9 | 10 | 11 | 12 | 13 | 14 | 15 | 16 | 17 | 18 | 19 | 20 |

191 kDa
97 kDa
64 kDa


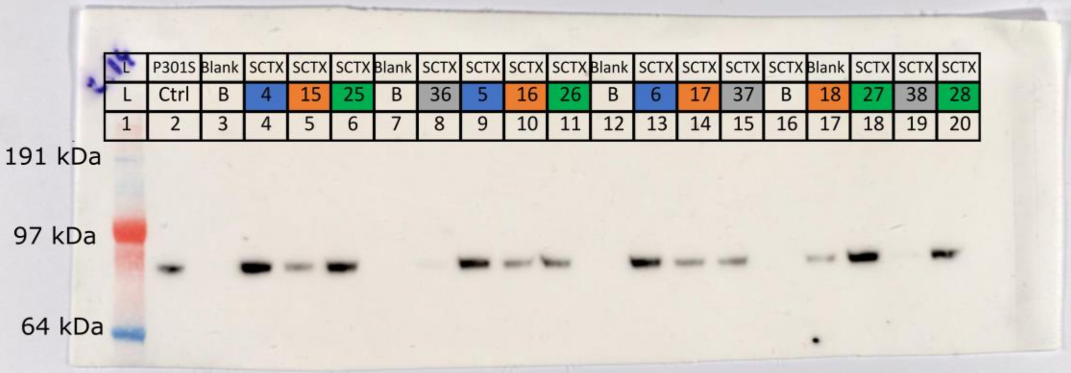

| L | P301S | Blank | SCTX | SCTX | SCTX | Blank | SCTX | SCTX | SCTX | SCTX | Blank | SCTX | SCTX | SCTX | SCTX | Blank | SCTX | SCTX | SCTX |
|---|---|---|---|---|---|---|---|---|---|---|---|---|---|---|---|---|---|---|---|
| L | Ctrl | B | 4 | 15 | 25 | B | 36 | 5 | 16 | 26 | B | 6 | 17 | 37 | B | 18 | 27 | 38 | 28 |
| 1 | 2 | 3 | 4 | 5 | 6 | 7 | 8 | 9 | 10 | 11 | 12 | 13 | 14 | 15 | 16 | 17 | 18 | 19 | 20 |

191 kDa
97 kDa
64 kDa


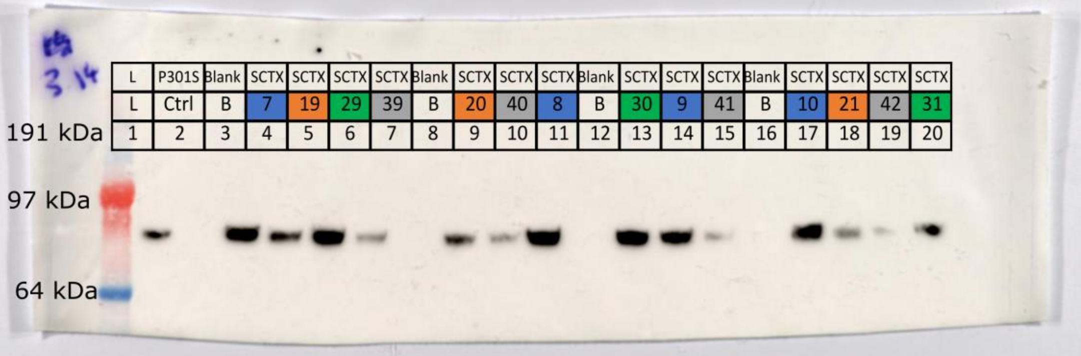

3.14

| L | P301S | Blank | SCTX | SCTX | SCTX | SCTX | Blank | SCTX | SCTX | SCTX | Blank | SCTX | SCTX | SCTX | Blank | SCTX | SCTX | SCTX | SCTX |
|---|---|---|---|---|---|---|---|---|---|---|---|---|---|---|---|---|---|---|---|
| L | Ctrl | B | 7 | 19 | 29 | 39 | B | 20 | 40 | 8 | B | 30 | 9 | 41 | B | 10 | 21 | 42 | 31 |
| 1 | 2 | 3 | 4 | 5 | 6 | 7 | 8 | 9 | 10 | 11 | 12 | 13 | 14 | 15 | 16 | 17 | 18 | 19 | 20 |

191 kDa
97 kDa
64 kDa

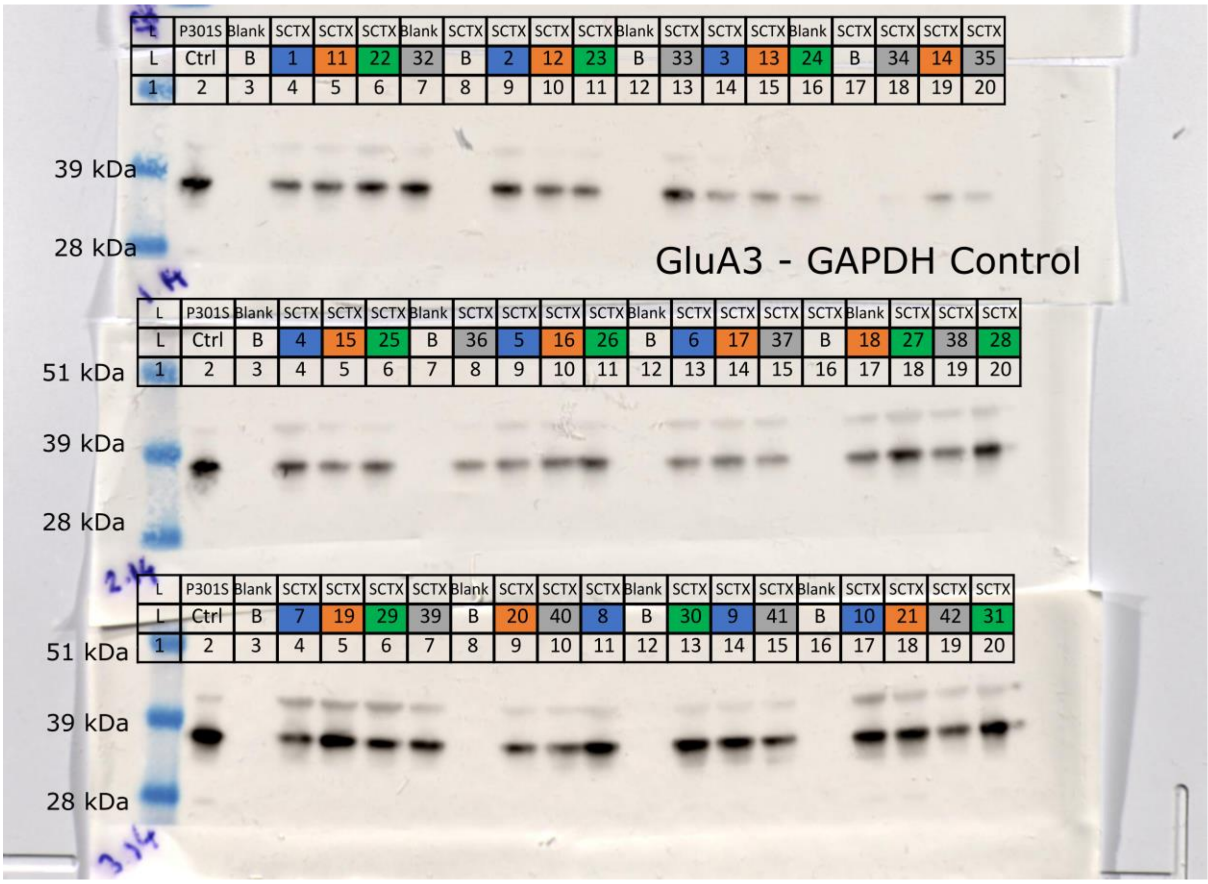
L P301S Blank SCTX SCTX SCTX Blank SCTX SCTX SCTX SCTX Blank SCTX SCTX SCTX Blank SCTX SCTX SCTX SCTX
L Ctrl B 1 11 22 32 B 2 12 23 B 33 3 13 24 B 34 14 35
1 2 3 4 5 6 7 8 9 10 11 12 13 14 15 16 17 18 19 20
39 kDa
28 kDa
GluA3 - GAPDH Control
L P301S Blank SCTX SCTX SCTX Blank SCTX SCTX SCTX SCTX Blank SCTX SCTX SCTX SCTX Blank SCTX SCTX SCTX
L Ctrl B 4 15 25 B 36 5 16 26 B 6 17 37 B 18 27 38 28
1 2 3 4 5 6 7 8 9 10 11 12 13 14 15 16 17 18 19 20
51 kDa
39 kDa
28 kDa
L P301S Blank SCTX SCTX SCTX SCTX Blank SCTX SCTX SCTX Blank SCTX SCTX SCTX Blank SCTX SCTX SCTX SCTX
L Ctrl B 7 19 29 39 B 20 40 8 B 30 9 41 B 10 21 42 31
1 2 3 4 5 6 7 8 9 10 11 12 13 14 15 16 17 18 19 20
51 kDa
39 kDa
28 kDa

### GluN1

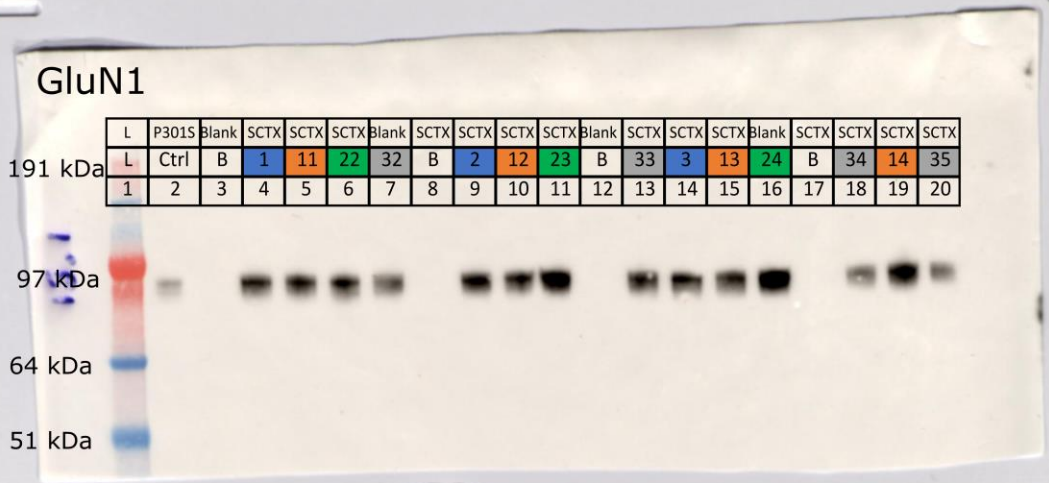

L P301S Blank SCTX SCTX SCTX Blank SCTX SCTX SCTX SCTX Blank SCTX SCTX SCTX Blank SCTX SCTX SCTX SCTX
L Ctrl B 1 11 22 32 B 2 12 23 B 33 3 13 24 B 34 14 35
1 2 3 4 5 6 7 8 9 10 11 12 13 14 15 16 17 18 19 20
191 kDa
97 kDa
64 kDa
51 kDa


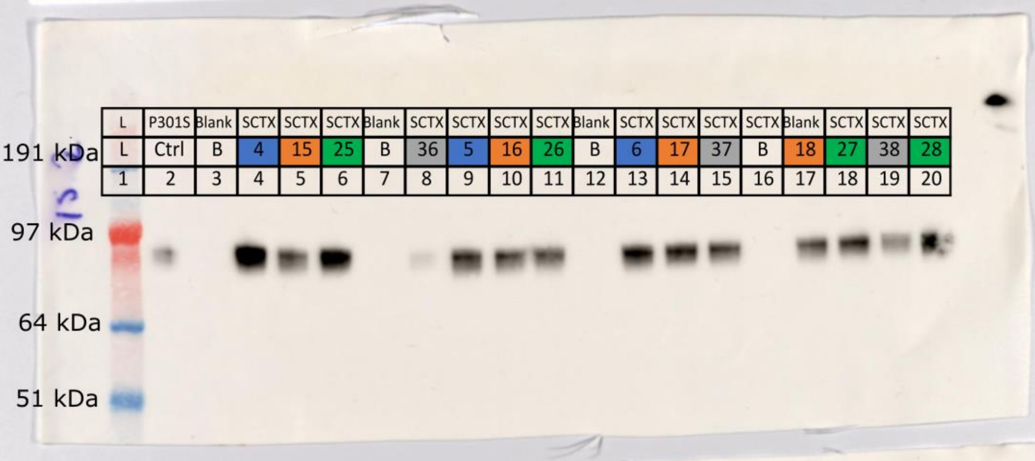

L P301S Blank SCTX SCTX SCTX Blank SCTX SCTX SCTX SCTX Blank SCTX SCTX SCTX SCTX Blank SCTX SCTX SCTX
L Ctrl B 4 15 25 B 36 5 16 26 B 6 17 37 B 18 27 38 28
1 2 3 4 5 6 7 8 9 10 11 12 13 14 15 16 17 18 19 20
191 kDa
97 kDa
64 kDa
51 kDa


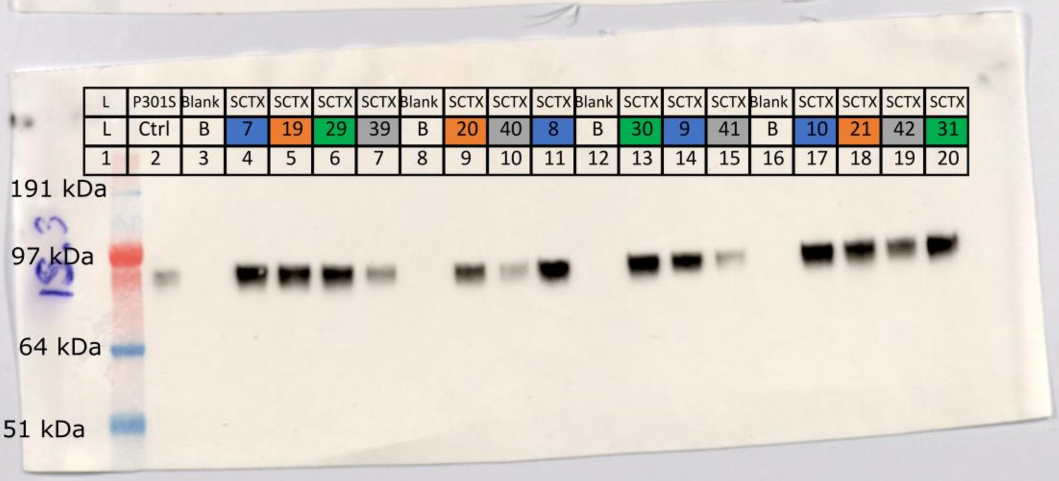

L P301S Blank SCTX SCTX SCTX SCTX Blank SCTX SCTX SCTX Blank SCTX SCTX SCTX Blank SCTX SCTX SCTX SCTX
L Ctrl B 7 19 29 39 B 20 40 8 B 30 9 41 B 10 21 42 31
1 2 3 4 5 6 7 8 9 10 11 12 13 14 15 16 17 18 19 20
191 kDa
97 kDa
64 kDa
51 kDa

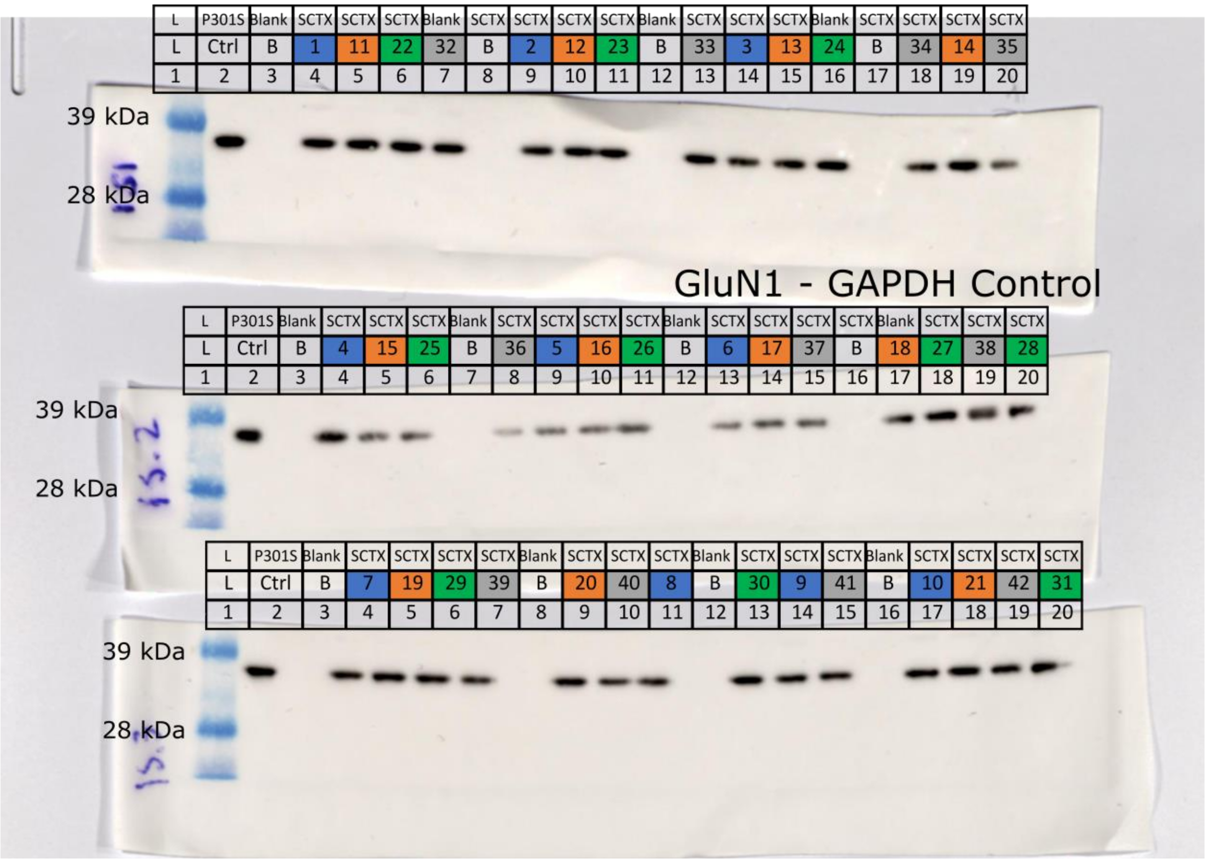


GluN1 - GAPDH Control
